# A CROSS-SPECIES ANALYSIS OF CELL WALL MECHANOSENSORS

**DOI:** 10.1101/2025.11.24.690177

**Authors:** Celia Municio-Diaz, Nicolas Minc

**Author notes:** Correspondence to NM.

## Abstract

The cell wall (CW) protects fungal cells from various mechanical challenges making its integrity essential for cell survival. CW integrity is monitored by transmembrane sensors that activate downstream effectors to promote CW synthesis in response to injuries. Sensors of the WSC family are found in most fungi, and share a conserved architecture, with a cytoplasmic tail, a single transmembrane domain and a long Serine Threonine Rich domain (STR) prolonged by a WSC domain, both embedded in the CW. In response to forces applied onto the CW, these extracellular domains promote force detection, sensor clustering and cell survival. Interestingly, Wsc sensors exhibit variations in domain sequence and size among different fungal species. To understand how these variations impact mechanosensing, we heterologously expressed Wsc sensors taken from *S. cerevisiae* and *C. albicans*, in the fission yeast *S. pombe*. Remarkably, we found that a subset of these foreign sensors could cluster at sites of CW compression, but that others failed, suggesting divergences in mechanosensing abilities. By swapping sensor domains, we demonstrate that both the cytoplasmic tail and STR influence mechanosensation. These findings reveal a high level of functional plasticity in fungal sensors, and identify tuneable modules that may regulate mechanosensing of various CWs.

**SIGNIFICANCE STATEMENT:**

- Cell Wall Mechanosensors of the WSC family are present in most fungi, but whether they can detect mechanical stimuli in a foreign cell wall of a distant fungal species is unknown.
- The authors heterologously expressed Wsc sensors taken from *S. cerevisiae* and *C. albicans* in the fission yeast *S. pombe* and demonstrate that a subset of foreign sensors can probe mechanical stress in its Cell Wall.
- This work highlights a remarkable plasticity in mechanosensors ability to detect mechanical stress in the Cell Wall and identifies domains involved in regulating mechanosensing.

## INTRODUCTION

Surface mechanosensing is of fundamental importance for cellular physiology, as it allows cells to adapt processes such as adhesion, motility or morphology to the mechanical properties of their environment (Wang *et al*., 1993; Wozniak and Chen, 2009). Walled cells, such as those of fungi and plants, are generally non-motile and non-adherent, but they make use of surface mechanosensing to survey the integrity of their protective Cell Wall (CW). Indeed, these cells feature cytoplasmic turgor pressures of few tens of atmospheres that put CWs under unusually large tensional stresses, causing risks of CW failure and consequent cell lysis (Mishra *et al*., 2022; Municio-Diaz *et al*., 2022). These risks are mitigated by the Cell Wall Integrity pathway (CWI), a signalling cascade that monitors and regulates CW integrity (Perez and Cansado, 2010; Levin, 2011). This cascade starts with upstream transmembrane sensors directly embedded into the CW. These sensors detect mechanical injuries or excessive stress/strain in the CW to activate downstream effectors including the RHO GTPase Rho1 that promote CW synthesis, as well as a MAPK module that turns on the expression of a set of specific CW regulatory genes (Perez and Cansado, 2010; Levin, 2011). In general, however, the mechanisms by which sensors detect mechanical signals remain poorly understood.

In fungi, surface sensors belong to two conserved families: the WSC (Wall Surface Component) and MID (Mating Induced Death) families. Depending on species, these sensors may be essential for cell survival, or only promote CW integrity during certain life instances, such as mating (Perez and Cansado, 2010; Rodicio and Heinisch, 2010; Levin, 2011). The fission yeast *Schizosaccharomyces pombe*, features for instance 2 main sensors : the WSC-type Wsc1 and the MID-type Mtl2 which complement each other to support cell viability (Cruz *et al*., 2013). Sensors from these two families share a similar architecture, with a C-ter cytoplasmic tail that communicates with downstream effectors, a single-pass transmembrane domain and a long STR (Serine Threonine Rich) domain that protrudes into the CW. Sensors of the WSC families possess a CRD (Cystein Rich Domain, also called WSC domain) at their N-ter, while sensors of the MID family only feature an N-glycosylated Asp residue (Hutzler *et al*., 2008). The STR domain is thought to play a major role in mechanosensing. First, this domain is highly mannosylated, allowing it to bind CW polysaccharides (Elhasi and Blomberg, 2019). Second, it has been proposed to adopt a “nano-spring” configuration, plausibly deforming along with the CW to sense surface stresses or strains and activate the CWI (Dupres *et al*., 2009). Recent studies in the fission yeast *S. pombe*, have shown that both STR and WSC domains in the Wsc1 sensor promote the formation of micrometric clusters at sites of CW mechanical compression; providing a direct visual output for surface mechanosensing (Neeli-Venkata *et al*., 2021).

Although they exhibit a general conserved architecture, sensors vary in the specific sequence of their domains among fungal species. In addition, certain species, like budding yeast for instance, feature several members of the Wsc or Mid sensors families, that support CW integrity in different contexts (e.g. mitotic growth vs mating) (Levin, 2011). Especially, the STR domain, is generally built from disordered ST repeats, but can exhibit large variations in sizes, as well as in the fraction of S and T amino acids, that serve as sites for O-glycosylation (Kock *et al*., 2015). These considerations raise the interesting possibility that sensor size or sequence may be adapted to probe different type or amplitude of mechanical stress among different fungal CWs that vary in thickness, composition, or turnover. To date, however, there has been no attempt to directly compare the behaviour and function of diverse CW sensors.

Here, using fission yeast as a model system, we heterologously expressed five sensors taken from *S. cerevisae* and *C. albicans*, which diverge from *S. pombe* by hundreds of Myrs (Sipiczki, 2000). We found that some sensors exhibited a relatively diffuse localization pattern reminiscent of the native Wsc1sp, while others were more polarized. Strikingly, some of these sensors could detect local compressive forces applied on the CW, by forming local clusters like the native *S. pombe* Wsc1sp. By generating swapped domains alleles, we could attribute localization patterns to C-ter tails which recruit the endocytic machinery, and variations in mechanosensing abilities to both C-ter tails and STR domains. Therefore, this work identifies modules and domains that may tune CW mechanosensing among fungi.

## RESULTS

### Heterologous expression and localization of Wsc sensors in *S. pombe*

WSC-like surface sensors can be found in fungal species across all the Ascomycota phylum. These species exhibit various morphologies, lifestyles and CW compositions, raising the interesting hypothesis that some specific features of WSC sensors could have evolved to optimize the detection of particular mechanical challenges or CW compositions. In order to explore how sensor sequences or sizes may impact their mechanosensing capacities and function, we selected 4 WSC-type sensors from *S. cerevisiae*, and 1 from *C. albicans.* Importantly, the CWs of *S. cerevisiae* and *C. albicans* differ in thickness and composition with that of *S. pombe*, notably featuring chitin as an important CW polymer, that is absent in *S. pombe* CW (Perez and Ribas, 2004). The selected foreign sensors covered a range of STR size from 115 aa to 304 aa similar to the range found across fungi (Fig 1A). The selected *S. cerevisiae* sensors are Slg1sc, Wsc2sc, Wsc3sc and Wsc4sc. They have been characterized to support CW integrity during multiple stress conditions (Verna *et al*., 1997; Zu *et al*., 2001; Vélez-Segarra *et al*., 2020). The *C. albicans* sensor Wsc1ca is smaller than other sensors, and was also previously shown to affect CW regulation (Norice *et al*., 2007) . These five sensors exhibited ∼27-36% in protein sequence identity with the native Wsc1sp, with Wsc2sc and Wsc3sc being the closet and Slg1sc, Wsc4sc and Wsc1ca, the most distant (Fig 1B). Domain sequence similarities analysis suggested a relatively large diversity in most protein domains, with the largest variability at the level of the cytoplasm C-terminal domain (Fig 1C).

**Figure 1.**
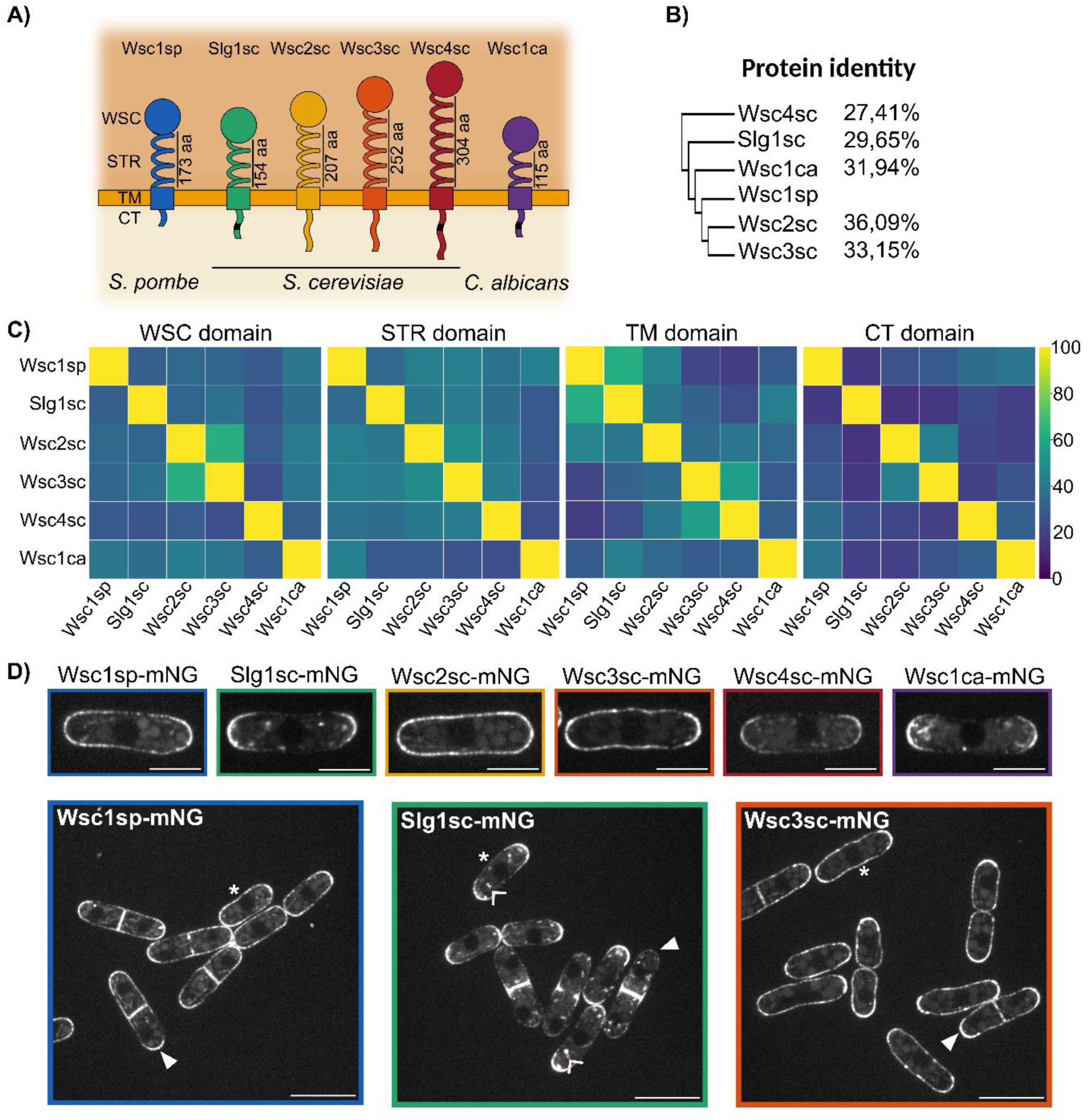
Heterologous expression of Wsc homologous sensors in *S. pombe*. A) Schematic models of sensor domains. WSC: cell Wall Stress-responsive Component domain; STR, Serine Threonine Rich domain; TM, TransMembrane domain; CT, Cytoplasmic Tail. Numbers correspond to the amino acid length of the STR. Black squares on the CT represent endocytic signals. B) Guide tree based on the full sequence similarity between sensors. The numbers are the percentage of amino acids that are identical at corresponding positions (identity) compared to *S. pombe* Wsc1sp. C) Heat maps showing the protein identity between sensors domains. D) Representative spinning disk mid-slice images showing the localization of the different tagged sensors, carrying the signal peptide of Wsc1sp, in agar pads. Upper row: Localization during interphase. Lower row: Characteristic localization during the cell cycle of the two sensors classes represented by Slg1sc, which is highly polarized, and Wsc3sc, which has a more homogeneous distribution. Asterisks indicate the side of the cells during interphase, filled arrows point the tips during septation and open arrows point to vesicles in the cytoplasm containing the sensor. Scale bars: individual cells 5 µm, group of cells 10 µm.

We generated *S. pombe* strains in which these 5 homologous genes were inserted in place of the native Wsc1sp allele, to ensure similar expression profile and transcriptional regulation. To allow for live visualization, these sensors were fused at their C-ter with a mNeonGreen fluorescent protein. All five sensors exhibited a pronounced localization to the plasma membrane, but some of them, like Wsc4sc, had less fluorescent signal on the surface and a higher signal in the cytoplasm (Fig S1A). To improve membrane targeting, we thus added a N-terminal signal peptide taken from Wsc1sp. This generally improved the fluorescent level at the cell surface, without altering the general localization of sensors without this native signal peptide (Fig 1D). Hence, all follow up constructs in this study feature this native Wsc1sp signal peptide to optimize surface targeting.

Interestingly, the localization of these five sensors was distinct, and could be separated into two general classes. Wsc2sc and Wsc3sc, had a relatively diffuse localization around cells, with some level of enrichment at cell tips in interphase, and at both the septum and cell tips during cell division. This localization pattern, was reminiscent of the localization pattern of the native Wsc1sp (Fig 1D and Fig S1B-C). In contrast, Slg1sc and Wsc1ca were highly enriched at cell tips, with almost no detectable signal on cell sides in interphase, and were completely re-targeted to the septum during cell division. In addition, these two sensors were also present in recycling vesicles in the cytoplasm (Fig 1D). This localization pattern, was similar to that of CW transmembrane enzymes like Bgs4 and Bgs1 (Cortes *et al*., 2002, 2005). Finally, Wsc4sc had a pattern of localization intermediate between these two classes, being more polarized than Wsc1sp, but to a lesser extent than Slg1sc and Wsc1ca (Fig 1D and Fig S1C). We conclude that foreign sensors can be expressed and localized in the plasma membrane and CWs of *S. pombe*, but that variability in their sequences causes them to be polarized along distinct patterns.

### Endocytic signals in C-ter cytoplasmic tails regulate sensor localization

The distinct localization patterns of the sensors suggested a plausible regulation by membrane recycling. Indeed, endocytosis has been shown to serve as a major module to promote the polarization of transmembrane proteins in yeast (Valdez-Taubas and Pelham, 2003). To explore this hypothesis, we searched for putative endocytic signals in the C-ter of these sensors. We could not identify any specific endocytic signal sequence in Wsc1sp, Wsc2sc and Wsc3sc, in agreement with their more diffuse localization around the cell surface. However, Slg1sc featured a NPFDD sequence, known to target proteins to the endocytic pathway. Similarly, Wsc4sc, featured a SPFHD sequence and Wsc1ca a NPFQHPADD, both pertaining to general consensus sequence that prime endocytic recycling (Piao *et al*., 2007). These results align with the presence of these 3 sensors in recycling cytoplasmic vesicles. They suggest that the more polarized localization of Slg1sc, Wsc4sc and Wsc1ca may emerge from the presence of an endocytic signal at their C-ter.

To directly test the role of endocytosis, we first expressed these sensors in a *end4Δ* mutant, previously shown to be highly defective in endocytosis (Iwaki *et al*., 2004). Strikingly, all five sensors expressed in this background, now exhibited a more diffuse signal similar to that of the native Wsc1sp (Fig 2A-B). Second, we generated swapped alleles by replacing the C-ter tail of all foreign sensors with that of the native Wsc1sp. The C-ter of Wsc1sp does not feature any obvious signal for endocytosis, and its deletion does not grossly affect its localization (Neeli-Venkata *et al*., 2021). Slg1sc, Wsc4sc and Wsc1ca chimeric sensors with Wsc1sp C-ter tail, now exhibited a loss of polarization, with a clear signal along cell sides in interphase and retention of tip localization during cell division (Fig 2C-2D). This localization pattern was similar to that of Wsc1sp, directly attributing their polarized distribution to the presence of endocytic signals in their C-ter tails. Surprisingly, however, we found that swapping the C-ter of Wsc1sp into Wsc2sc, caused it to become highly polarized. Together these results demonstrate that endocytic recycling of some of these sensors, promoted by specific consensus signals in their C-ter cytoplasmic tails, may serve as a powerful module to polarize them to growing tips and septa.

**Figure 2.**
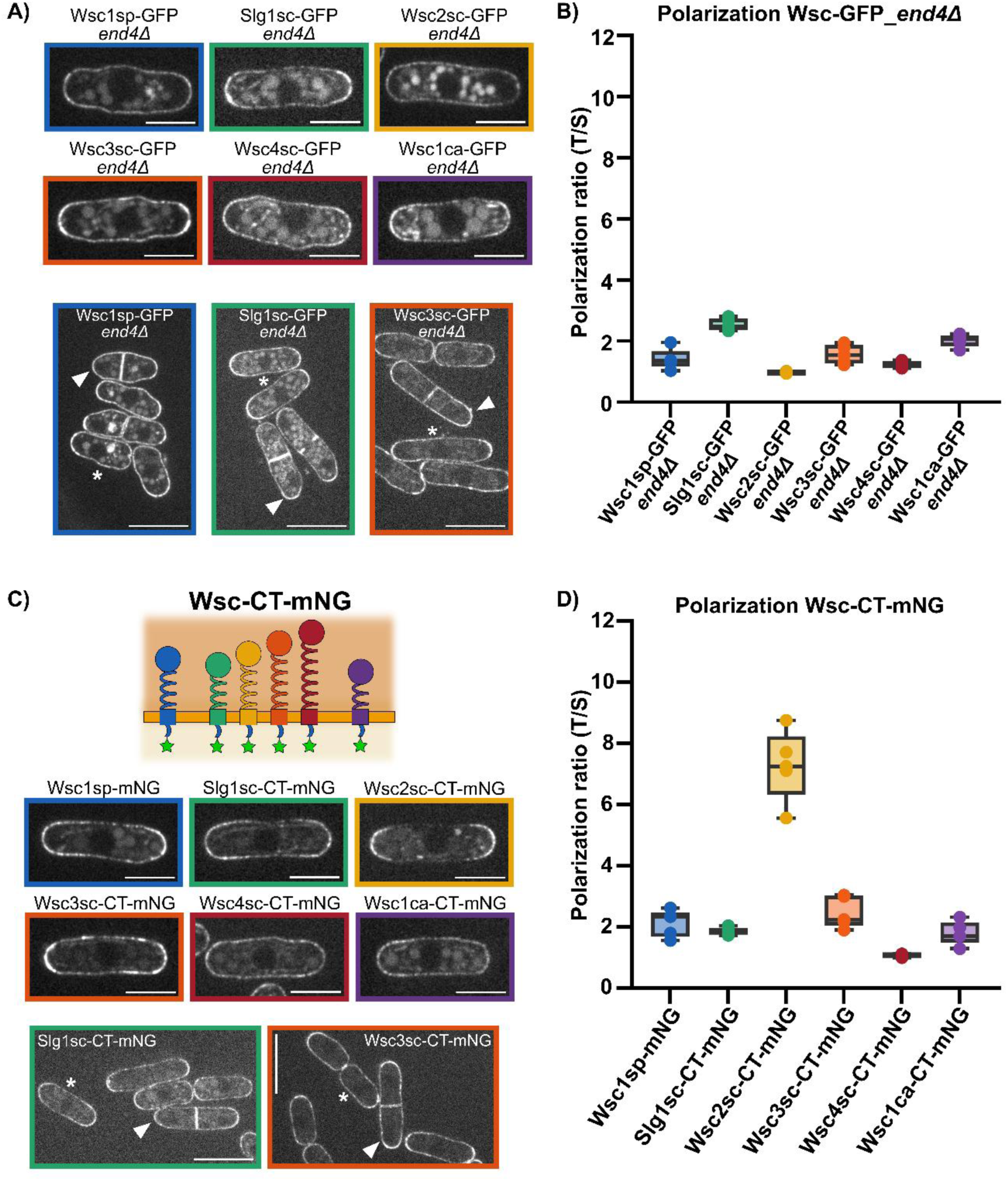
Endocytic signals in the C-terminal cytoplasmic tail influence sensor localization. **A)** Representative images showing the localization of the different tagged sensors in an *end4Δ* mutant. Upper rows: Localization during interphase. Lower row: Localization during the cell cycle. **B)** Polarization of the sensors in *end4Δ* mutants. This ratio is calculated as the signal intensity in the tip over the signal on the sides. Values close to 1 indicate a homogeneous distribution of the sensor, > 1 polarization of the sensor and < 1 accumulation at cell sides. **C)** Schematic of sensors in which the CT has been swapped with that of Wsc1sp. Localization of the indicated sensors. Upper rows: Localization during interphase. Lower row: Localization during the cell cycle. **D)** Polarization of the CT swapped sensors. In A) and C), asterisks indicate the side of the cells during interphase, filled arrows point to the tips during septation. Scale bars: individual cells image 5 µm, group of cells images 10 µm. In B) and D) n = 5 cells for each sensor.

### Mechanosensation of *S. pombe* Cell Wall deformation by foreign sensors

We previously showed that Wsc1sp exhibits a clustering behaviour at local sites where CWs are compressed, either by the presence of a neighbouring cell or an inert physical barrier. Clustering behaviour happened in dose-dependence with applied stress on CWs, and was reversed when stress was released (Neeli-Venkata *et al*., 2021). Among assays used to characterize this behaviour, we had developed linear microfabricated channels, in which *S. pombe* cells grow and press onto each other, recruiting Wsc1p at sites of compressed cell-cell contacts (Neeli-Venkata *et al*., 2021). Therefore, to assay mechanosensing capacities of foreign sensors in *S. pombe*, we grew cells expressing each sensor in microchannels until they reached a high degree of crowding characterized by triangular-like cell shapes and compressed cell-cell contact, detectable as flat interfaces between cells (Fig 3A) (Haupt *et al*., 2018; Neeli-Venkata *et al*., 2021). We quantified the enrichment of sensors to these compressed sites and compared them to their enrichment at free cell tips at channel ends, and normalized by a factor 2 to account for the presence of 2 juxtaposed membranes at cell-cell contacts (Fig 3A-3B).

**Figure 3.**
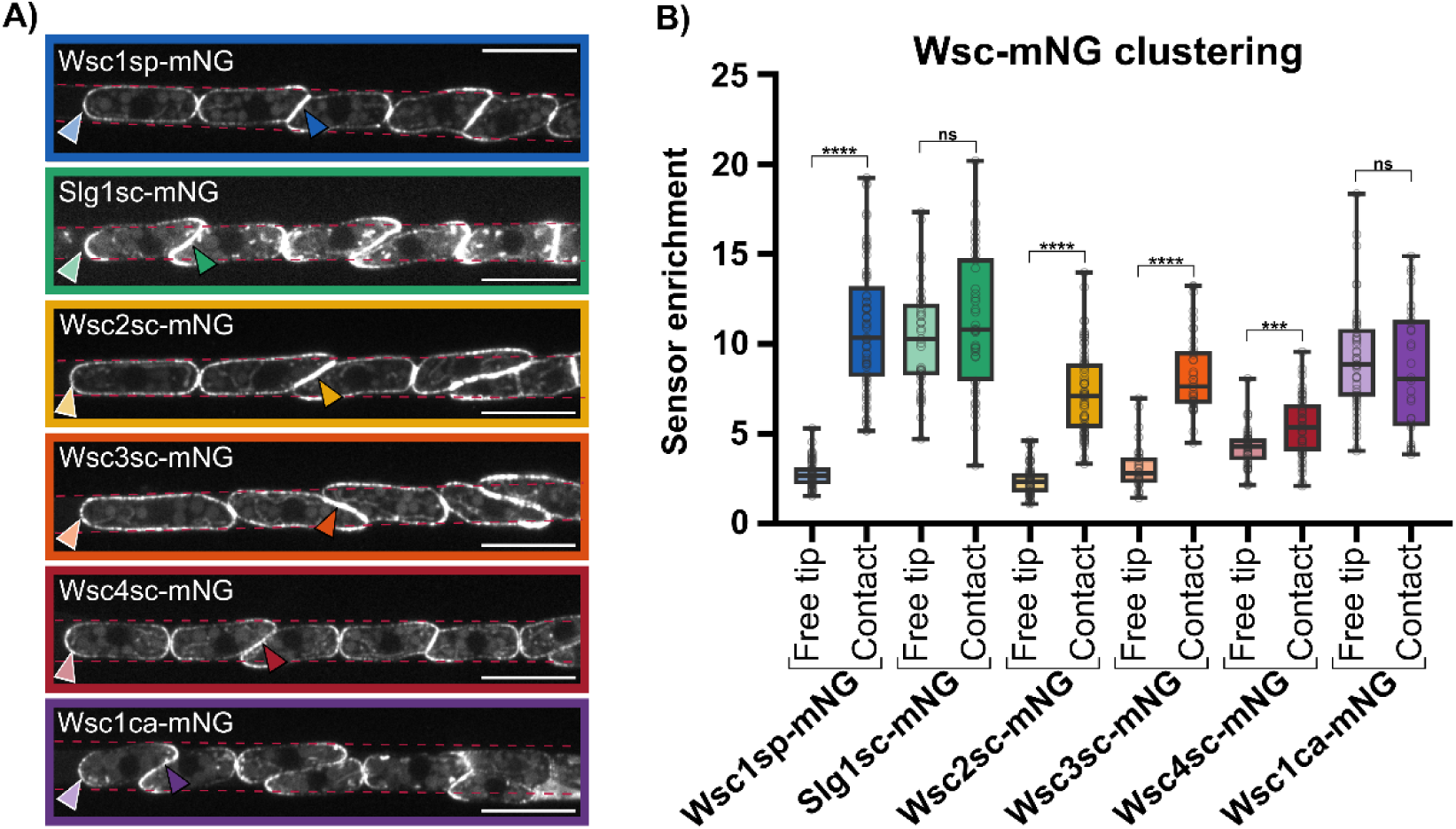
Sensor clustering under mechanical stress. **A)** Representative images of *S. pombe* cells expressing the sensors grown in linear microchannels up to a high confinement regime. Light and dark-colored arrows point to examples of the quantified free tips and pressurized cell-cell contacts respectively. Dashed red lines indicate the microchannels walls. **B)** Box-plot showing the enrichment at free tips and at cell contacts for the different sensors tagged with mNeonGreen. n= 54, 61, 59, 58, 61, 60, 58, 69, 29, 32, 48, 53 cells, respectively. Comparison between free tips and contacts was done using the Mann-Whitney test, T-test P-values, **, P < 0.01 ***, P < 0.001, ****, P < 0.0001. Scale bars, 10 µm.

Remarkably, Wsc2sc and Wsc3sc, exhibited a strong enrichment at sites of CW compression, forming clusters comparable in size and intensity as those formed by the native Wsc1sp. Wsc4sc was also slightly enriched at sites of compressed CWs as compared to free growing tips, but to a lesser extent than Wsc2sc and Wsc3sc (Fig 3A-3B). In contrast, Slg1sc and Wsc1ca did not exhibit any specific enrichment at sites of cell contacts, beyond what their strong tip polarization pattern could yield (Fig 3A-3B). These results demonstrate that built-in properties of some WSC sensors allow them to detect enhanced CW stress or strain, even in a foreign organism with a different CW architecture and composition.

To test if the presence of endocytic signals in Slg1sc, Wsc4sc and Wsc1ca could alter mechanosensing, we assayed their clustering in an *end4Δ* background or used chimeric sensors with a Wsc1sp C-ter tail to redistribute them around cells. When grown in microchannels, these three sensors that could not be recycled, now exhibited a mild but significant enrichment at sites of CW compression, that was nonetheless smaller than that of the native Wsc1sp (Fig S2A-D). This suggests that sensor endocytosis, mediated by their C-ter tails, can have a significant negative impact on mechanosensation capacities.

### Impact of extracellular domains on mechanosensing

We next sought to test if and how extracellular STR and WSC domains may impact localization and/or mechanosensation of foreign sensors. To explore the contribution of these domains, we generated swapped alleles that contain all domains of the native *S. pombe* Wsc1sp but in which we replaced the WSC or STR domain with that of the different foreign sensors (Fig 4B and 4F).

**Figure 4.**
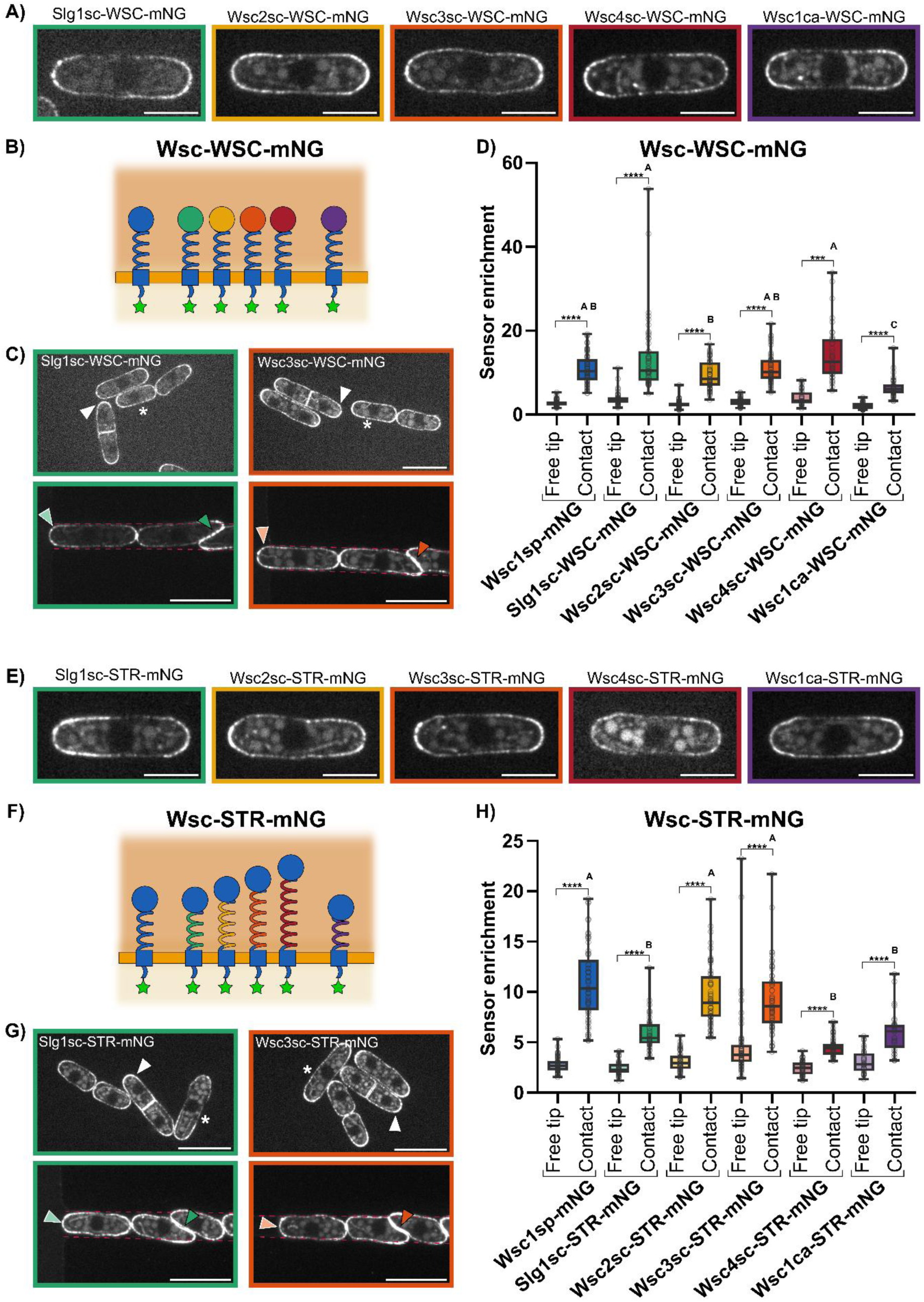
Role of the WSC and STR extracellular domains on sensor clustering under mechanical stress. **A) and E)** Representative images showing the localization of the Wsc-WSC-mNG and Wsc-STR-mNG sensors during interphase. Scale bars 5 µm. **B) and F)** Schematic representation of the Wsc-WSC-mNG and Wsc-STR-mNG sensors. In these swapped alleles, the WSC or the STR domain of the native Wsc1sp has been swapped with the WSC or the STR domain of the foreign sensors. **C) and G)** Characteristic images showing the localization of the sensors in agar pads (upper row) and in microchannels (lower row). White asterisks indicate the side of the cells during interphase, white arrows point to the tips during septation. Light and dark-colored arrows point to free tips and cell-cell contacts respectively. Scale bars 10 µm. **D) and H)** Boxplot showing the enrichment of the Wsc-WSC-mNG and Wsc-STR-mNG sensors at free tips and at cell contacts. In D) n = 54, 61, 46, 51, 51, 58, 48, 53, 35, 37, 39 and 40 cells and in H) n = 54, 61, 42, 42, 54, 61, 60, 60, 52, 50, 35 and 32 cells. The comparison between free tips and contacts was done with the Mann-Whitney test, T-test P-values, ***, P < 0.001, ****, P < 0.0001. The letters define groups which are significantly different from other groups after comparing the sensors enrichment at cell contacts, using the Kruskal-Wallis and Dunn’s tests.

In general, the WSC domains of these foreign sensors, exhibit a weak sequence similarity with that of Wsc1sp, but they always feature a conserved sets of cysteines thought to promote the proper folding and function of this domain (Fig 1B) (Schöppner *et al*., 2022). Accordingly, WSC swapped alleles did not exhibit any major changes in terms of localization and in clustering behaviour in microchannels, as compared to the native Wsc1sp. Yet, we noted a minor reduction in enrichment at cell contacts for the swapped Wsc1ca-WSC, and a minor improvement for the swapped Slg1sc-WSC and Wsc4sc-WSC (Fig 4A-D). Therefore, differences in WSC domain sequence among sensors, may only have a minor impact on mechanosensation capacities.

STR domains from different sensors also exhibit weak sequence similarities, and also feature major divergences in terms of length and content of S and T amino acids that serve as sites for O-mannosylation. The localization of swapped STR sensors was similar to that of the native Wsc1sp, indicating that STR domains do not influence protein localization (Fig 4E and G). However, their ability to sense compressive forces varied more. Wsc2sc-STR and Wsc3sc-STR exhibited an enrichment similar to the native Wsc1sp, suggesting that their STR has similar built-in properties for mechanosensation in spite of some degree of sequence divergence (Fig 4H). In contrast, Slg1sc-STR, Wsc1ca-STR, and Wsc4sc-STR were still significantly enriched, but to a level lower than for the native Wsc1sp. This suggests that their STR is partially defective in detecting forces applied on the *S. pombe* CW (Fig 4H). Together, these results indicate that STR domains play an important role in titrating mechanosensing abilities of different sensors.

### STR size and ST content do not influence mechanosensing

One major divergence among the 5 sensors selected comes from the size of their STR domains. Wsc2sc and Wsc3sc STRs are 207 aa and 252 aa long, quite comparable to the length of Wsc1sp STR which is 173 aa long, while Wsc1ca and Slg1sc have the shortest STR (115 aa and 154 aa, respectively) and Wsc4sc the longest (304 aa). Given previously proposed “nano-spring” models for STR based mechanosensing (Dupres *et al*., 2009), we reasoned that a plausible explanation for different degrees of mechanosensing could come from the size of STR domains as compared to the thickness of CWs. In that view, longer STRs like that of Wsc4sc could be better adapted to probe thicker CWs, while shorter ones like that of Wsc1ca thinner CWs.

To test this hypothesis, we first manipulated the length of STRs to shorten Wsc4sc-STR or lengthen Wsc1ca-STR and bring both of them to sizes similar to that of the native Wsc1sp STR (Fig 5A). This was achieved by removing the 149 central aminoacids of the STR domain for Wsc4sc and by duplicating a region of 40 aminoacids of the STR domain for Wsc1ca. However, we found no major changes in sensor localization or enrichment at cell-cell contacts, in spite of these drastic changes in STR length (Fig 5B). Second, we expressed Wsc1sp, Wsc1ca-STR and Wsc4sc-STR, that possess respectively the shortest and longest STR in a *kin1Δ* mutant (Fig 5C). This mutant exhibits a CW typically twice thicker than in WT cells (Cadou *et al*., 2010; Davi *et al*., 2019). In microchannels, the clustering of Wsc1sp was slightly reduced in *kin1Δ* cells, as compared to WT. However, the behaviour of both the long Wsc4sc-STR and the short Wsc1ca-STR was unaltered as compared to the WT background (Fig 5D). Altogether, these data strongly suggest that the length of STR domains related to CW thickness does not play any major role in mechanosensing.

**Figure 5.**
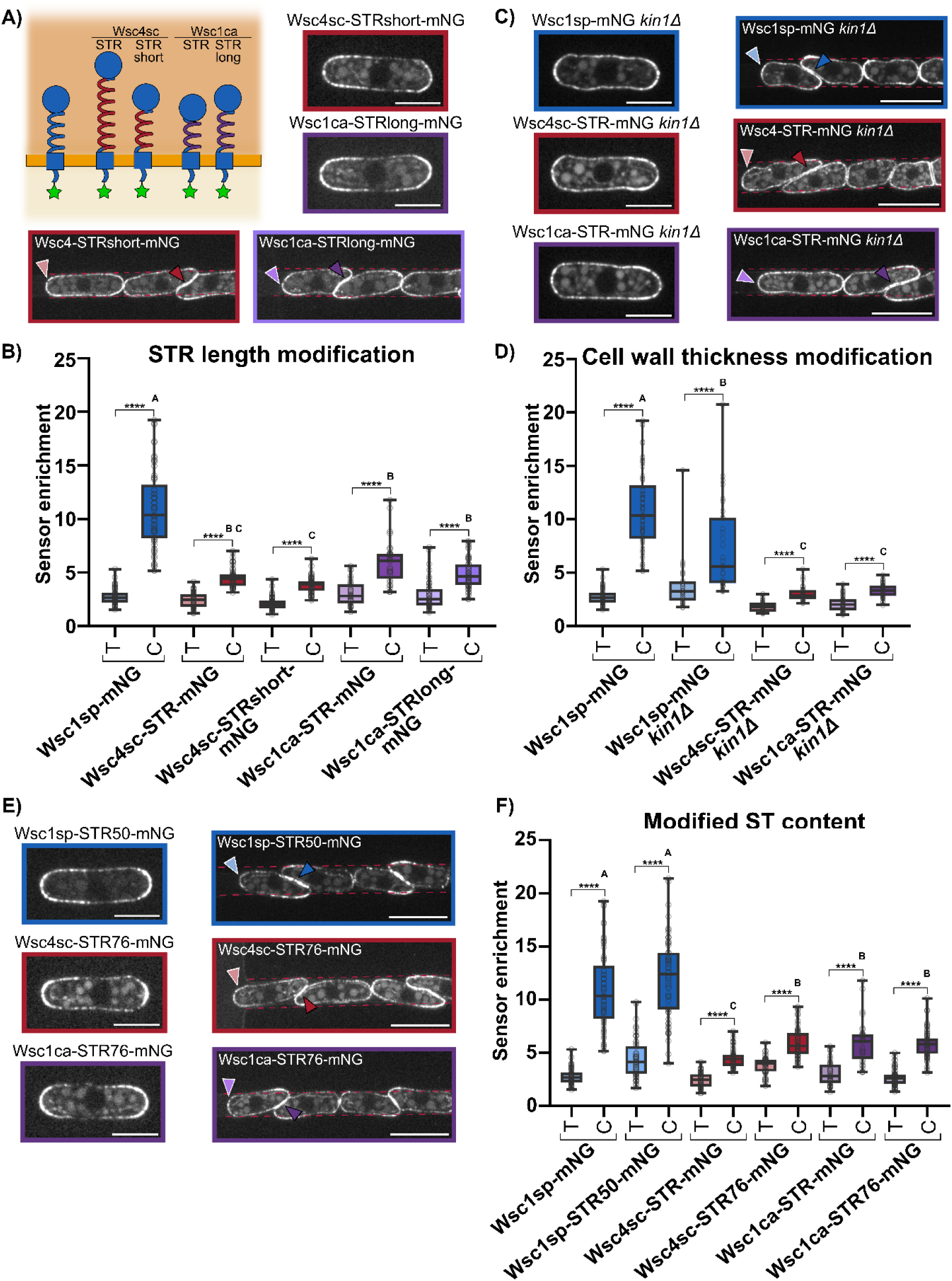
STR modifications in terms of length and in serine/threonine content does not influence sensors localization or clustering under force. **A)** Schematic showing the modifications in length of the STR domains for Wsc4sc and Wsc1ca. To have the same size as Wsc1sp STR (173 aa), Wsc4sc STR (304 aa) was shortened to 173 aa (STRshort) and Wsc1ca STR (115 aa) was lengthened to 173 aa (STRlong). Images showing the localization of the sensors during interphase and in microchannels. **B)** Enrichment of the sensors with modified STR length at free tips (T) and at cell contacts (C). n= 54, 61, 52, 50, 60, 59, 35, 32, 49 and 52 cells. **C)** Representative images showing the localization of the sensors in a *kin1Δ* mutant background during interphase and in microchannels. *kin1Δ* cells feature a CW twice as thick as in the WT. **D)** Enrichment of the sensors in a *kin1Δ* mutant background at free tips and at cell contacts. n = 54, 61, 32, 32, 33, 34, 50 and 41 cells. **E)** Representative images showing the localization of sensors in which the percentage of serines and threonines in the STR domain has been changed. In Wsc1sp-STR50-mNG the percentage of S and T has been reduced from 76 to 50%. In Wsc4sc-STR76-mNG and Wsc1ca-STR76-mNG this percentage has been increased from 61% and 56% respectively to 76 %. **F)** Enrichment of the sensors with modified S and T content at free tips and at cell contacts. n = 54, 61, 54, 60, 51, 57, 60 and 60 cells. In A), C) and E), light and dark-colored arrows point to free tips and cell contacts respectively. The comparison between free tips and contacts was done with the Mann-Whitney test, T-test P-values, ****, P < 0.0001. The letters define groups which are significantly different from other groups after comparing the sensors enrichment at cell contacts, using the Kruskal-Wallis and Dunn’s tests. Scale bars: individual cells 5 µm, channels 10 µm.

In addition to variation in STR length, we also noted a different proportion of Serine and Threonine residues in the STR domains of different foreign sensors (Fig 5E). Notably, Wsc1sp, Wsc2sc and Wsc3sc that exhibit the highest clustering behaviour, had a similar high degree of ST content of ∼60-70%. Conversely, Slg1sc, Wsc4sc and Wsc1ca which are defective in clustering feature a relatively lower degree of STs in their STRs below ∼60%. As these sites are thought to prime STR O-mannosylation and thus sensor binding efficiency to CW polymers, we sought to directly test their contribution to mechanosensing capacities. For this, we first reduced the proportion of STs from 76% to ∼50% in the native Wsc1sp STR by replacing a subset of S and T with alanine residues. However, we did not observe any difference in term of sensor clustering. Conversely, we increased the proportion of STs in the STRs of Wsc4sc and Wsc1ca to a proportion of ∼76% similar to that found in the STR of the native Wsc1sp. We noted no changes for Wsc1ca, and only a minor improvement in clustering behaviour for Wsc4sc. We note that although we expect these sequence changes to affect the degree of O-mannosylation of STRs, we could not directly assay how they modified O-mannosylation. Overall, these findings suggest, that the proportion of S and T in STR sequence, which presumably impact its degree of O-mannosylation does not grossly influence mechanosensing.

### Foreign sensor extracellular domains can support CW integrity

Finally, we sought to assay if foreign sensors, in spite of their divergence with the native Wsc1sp sensor, can support cell viability and CW integrity. For this, we used a *wsc1Δ* nmt1-Mtl2 shut off strain, which is not viable when supplemented with thiamine, given the absence of both Wsc1 and Mtl2 sensors to support CW integrity (Cruz *et al*., 2013). We expressed various foreign sensor alleles in this strain, and grew them in plates supplemented with thiamine to repress Mtl2 and assayed for viability. Expressing any of the full-length foreign sensors did not support survival, suggesting that these sensors cannot supplement the function of Wsc1sp in their native form. However, remarkably, expressing any of the five chimeric foreign sensors with the C-ter tail of the native Wsc1sp was sufficient to support cell viability. Similarly, both WSC and STR swaps, that contain the native C-ter tail also supported viability (Fig 6). Together, these data suggest that in spite of variations in mechanosensation abilities, all foreign sensor extracellular domains are sufficient to activate the downstream CWI, provided they are equipped with the correct C-ter tail for binding and activating downstream effectors. Importantly, however, although these data support that these sensors can support viability in normal growth conditions, we noted a significant amount of cell death for most of these alleles grown in microchannels, which were reminiscent of the behaviour of a *Wsc1Δ*. This suggests that the extracellular domains of foreign sensors may not completely support function in conditions of very high compressive stress on the CW (Fig S3).

**Figure 6.**
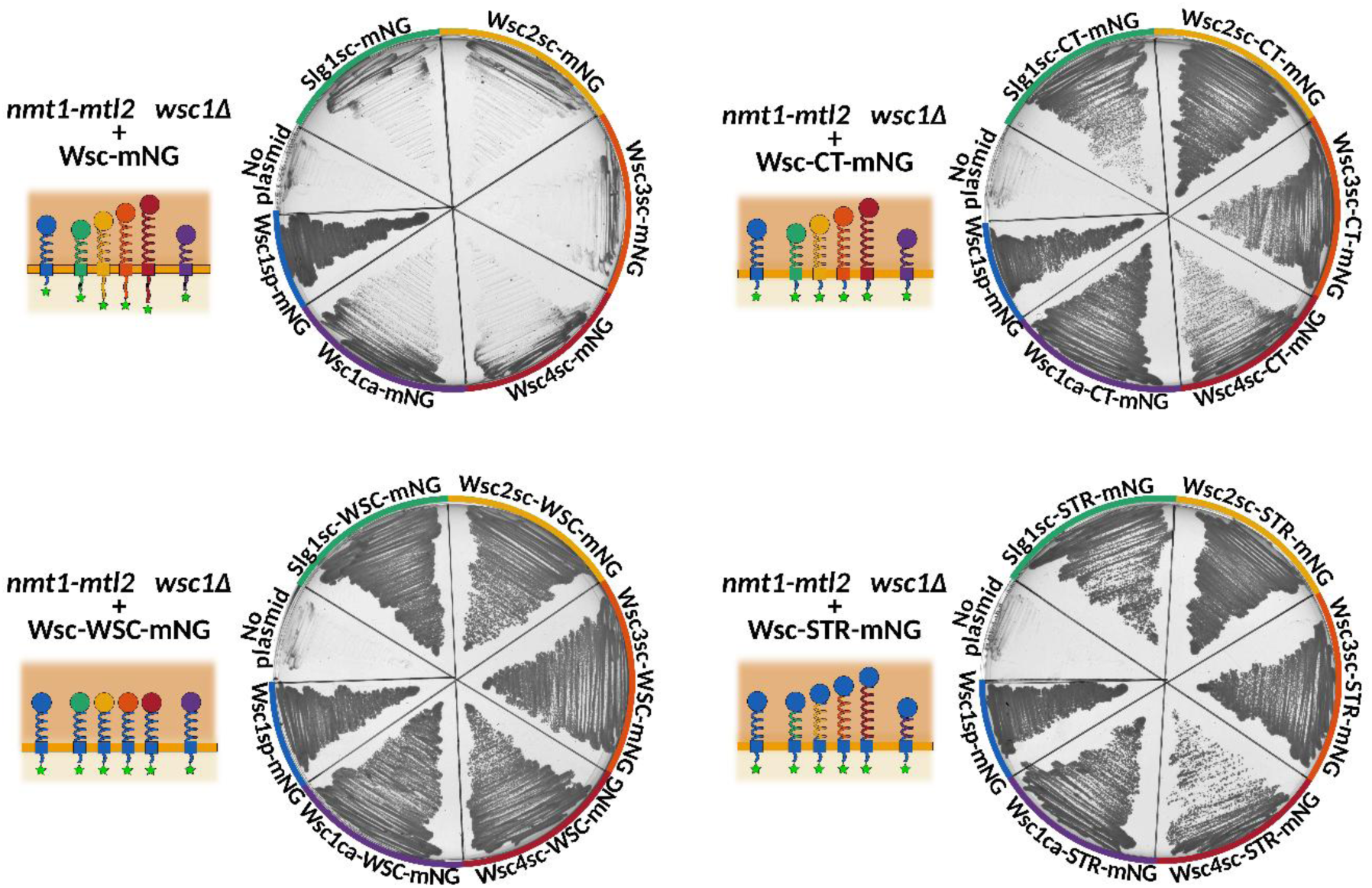
Functionality of foreign sensors in normal growth conditions. The strain nmt1-Mtl2 *wsc1Δ* is non-viable when grown on plates supplemented with thiamine due to the repression of Mtl2 under thiamine and therefore the absence of both sensors. The strain nmt1-Mtl2 *wsc1Δ* was transformed with plasmids bearing the indicated sensors and grown for 48h at 30°C in EMM-A agar plates supplemented with thiamine. No plasmid: Non transformed nmt1-Mtl2 *wsc1Δ* strain.

## DISCUSSION

Fungal cells are continuously exposed to surface mechanical stress caused by their high internal turgor, or the contact they form with growing neighbours in colonies. This makes CW mechanosensing a fundamental element for their survival and general behaviour. Interestingly, fungal hyphal cells display a large diversity in terms of tip growth speed or size, which are associated with drastic changes in turgor values, CW thickness or elasticity (Davi *et al*., 2019; Lemière and Chang, 2023; Chevalier *et al*., 2024). Therefore, it is plausible that mechanisms of mechanosensing may have been tuned during evolution to support particular levels of strains or stresses in the CW that vary depending on growth environment, values of turgor or CW properties.

Building on these considerations, we here assayed the behaviour of different CW mechanosensors of the WSC family coming from quite divergent fungal organisms in the foreign CW of *S. pombe* cells. A major finding of this work is to demonstrate that some foreign sensors that are presumably designed to function in specific CWs, can still properly localize and detect local forces applied onto a different CW in a quite distant organism. In addition, provided that they are equipped with the C-ter cytoplasmic tail of the native Wsc1sp sensor to interact with downstream effectors, these foreign sensors can support CW integrity and survival in the absence of other CW sensors in *S. pombe*. Such a functional plasticity suggests that some generic properties of STR and WSC domains are highly conserved in the WSC family allowing sensors to exert a basic function in most fungal CWs. We speculate that this could reflect that their extracellular domains interact with conserved CW polysaccharides, such as β- or α-Glucans which are highly abundant in most fungal CWs (Gow and Lenardon, 2023).

Another interesting observation of this study was to find that different sensors exhibit different degrees of clustering to sites of CW compressions, that we use as a proxy for mechanosensing abilities. Specifically, Wsc2sc and Wsc3sc exhibited a clustering behaviour, similar to the native Wsc1sp. In contrast, Wsc4sc had an intermediate clustering behaviour, and Slg1sc and Wsc1ca were completely defective in accumulating to sites of CW compression. By generating chimeric versions of these foreign sensors, we attributed these variations to changes in C-ter tail and STR sequences. Endocytic signals in the C-ter tail of Wsc4sc, Slg1sc, Wsc1ca sensors may promote their recycling thereby limiting their accumulation at sites of CW compression and thus their ability to detect local mechanical stimuli on the CW. Analysis and modulation of STR length or ST content did not reveal any specific changes, suggesting that other fine-tuned aspect of STR sequence such as the distribution of ST sites or mannosylation patterns could account for the various degrees of mechanosensing observed. Future comparative work of this kind will deepen our understanding for how evolution has modified specific protein sequences to support surface mechanosensation, and bares the promise of designing synthetic sensors optimized to detect particular level of mechanical stress or strain.

## MATERIAL AND METHODS

### Fission yeast strains and growth conditions

All strains used in this study are haploids, the genotypes and the source of the strains are listed in table S1. Standard *S. pombe* media and methods were used (Forsburg and Rhind, 2006). Genetic crosses were performed on malt extract agar plates. New generated strains in this study were obtained by tetrad dissection or random spore analysis and selection on appropriate media. To generate new strains via homologous recombination, a DNA fragment containing the Wsc1sp 3’UTR (∼1 k upstream the start codon), the sensor CDS, Wsc1sp 5’UTR (∼1k downstream the stop codon) and the resistance gene (ura4+) was obtained either by PCR (oCM9+oCM10) or by restriction enzymes from the plasmids mentioned below. Wild-type *S. pombe* strains were transformed by the lithium acetate method (Forsburg and Rhind, 2006). Correct integration of the transgene was analyzed by PCR with the oligonucleotide oCM83, in the 5’UTR region of *wsc1* outside the transgene, and a specific reverse primer inside the transgene (Table S2). For microscopy and microchannels experiments the strains were grown on liquid YE5S at 25°C.

### Cloning and DNA manipulations

DNA sequences were retrieved from PomBase (Rutherford *et al*., 2024) for *wsc1sp* (SPBC30B4.01c); from the Saccharomyces Genome Database for *slg1sc* (YOR008C), *wsc2sc* (YNL283C), *wsc3sc* (YOL105C) and *wcs4sc* (YHL028W); and from the *Candida* Genome Database for *wsc1ca* (C3_04250W_A). For expression in *S. pombe*, the genomic sequences of *slg1sc*, *wsc2sc*, *wsc3sc*, *wsc4sc* and *wsc1ca* were codon optimized for *S. pombe* and synthetized with a XbaI site at the 5’ and a NotI site at the 3’ (Eurofins Genomics, pCM1-pCM5). The length of the different domains of the sensors is listed in table S3.

All constructs were based on the plasmid pAL-*wsc1-GFP:ura4+* (pRZ21, Cruz et al., 2013) and are listed in table S4:

i. <u>Sensors with native signal peptide and C-terminally fused to GFP (SP-sensor-GFP):</u> The pRZ21 plasmid was modified by site directed mutagenesis to add a *NheI* site before the start codon of *wsc1sp* and remove the NotI site after the GFP but keeping the NotI site before the GFP (pCM10). The optimized and synthesized sensors sequences were then cloned between the NheI (compatible with XbaI overhangs) and NotI site (Plasmids pCM11-pCM15).
ii. <u>Sensors with *S. pombe* signal peptide and C-terminally fused to mNeonGreen</u> <u>(Sensor-mNG):</u> First, the DNA sequence corresponding to the 30 first amino acids of *S. pombe*, which contains the full signal peptide plus its cleavage site, was cloned in place of the full native signal peptide plus the first amino acid of the WSC domain to account for the native cleaving site of each of the other sensors (pCM16-pCM20). Second, the GFP sequence was replaced by the mNeonGreen sequence optimized for *S. pombe* (pCM52-pCM56) (a gift from Silke Hauf, Virginia tech). For Wsc1sp-mNG, the GFP of the pRZ21 plasmid was also replaced by mNeongreen (pCM60).
iii. <u>Sensors with the C-terminal tail of *S. pombe* and C-terminally fused to mNeonGreen</u> <u>(Sensor-CT-mNG)</u>: The plasmids pCM11-pCM15 were modified to replace the cytoplasmic tail of *S. cerevisiae* and *C. albicans* sensors by Wsc1sp cytoplasmic tail (DNA fragment corresponding to the amino acids 314 to 374) and exchange the GFP by m-NeonGreen (pCM35-pCM39).
iv. <u>Wsc1sp with the WSC domain from the other sensors and C-terminally fused to</u> <u>mNeonGreen (Wsc-WSC-mNG)</u>: The plasmids pCM35 and pCM39 were modified to replace the STR and transmembrane domains of Slg1sc (amino acids 111-285) and Wsc3sc (amino acids 133-405) by *S. pombe* Wsc1sp STR and transmembrane domains (amino acids 120-313). The resulting Wsc1sp sensors only contain the WSC domain of Slg1sc and Wsc3sc respectively and are fused to mNeonGreen (pCM44 and pCM45). The plasmid pCM44 was further modified to replace the WSC domain of Slg1sc by the WSC domains of Wsc2sc (amino acids 25-118), Wsc4sc (amino acids 28-110) and Wsc1ca (amino acids 29-116) (pCM57-pCM58).
v. <u>Wsc1sp with the STR domain from the other sensors and C-terminally fused to</u> <u>mNeonGreen (Wsc-STR-mNG):</u> The plasmid pCM60 was modified to replace the region of Wsc1sp between amino acids positions 128 and 282 by the STR domain of the other sensors (pCM61-pCM65).
vi. <u>Modification of the STR size of Wsc4sc and Wsc1ca:</u> To reduce the size of the STR domain of Wsc4sc, the sensor coded in the plasmid pCM64 was modified to remove the region between amino acids 187 and 335 (pCM67). To increase the size of Wsc1ca STR domain, the plasmid pCM65 was modified: the STR middle region (amino acids 160 to 199) was duplicated and inserted in position 199 (pCM166). In both cases, the resulting STR region, counted from Wsc1sp WSC to the transmembrane domain, has 173 amino acids like the one of *S. pombe*.
vii. <u>Modification of the serine and threonine content in Wsc1sp, Wsc4sc and Wsc1ca:</u> The STR region of Wsc1sp is highly glycosylated (Lodder *et al*., 1999; Lommel *et al*., 2004; Lõoke *et al*., 2011). O-glycosylation occurs in serine and threonine residues. In Wsc1sp, the STR domain contains a 64.74% and 11.56% of serines and theronines respectively (112 serines and 20 threonines in the 173 aa STR). The STR regions of Wsc4sc and Wsc1ca were modified to have the same length but increased S+T%, up to 76%, in the final STR coding region counting from WSC domain to transmembrane domain. For the modifications it was taken into account that in Wsc1sp the number of serines is higher than the threonines, therefore, the number of threonines was not modified but the number of serines was increased by replacing the other amino acids types. These changes aim to keep an overall proportion of amino acids similar to the one of Wsc1sp. The new STRs are named Wsc4STR76 and Wsc1caSTR76. For Wsc1sp, the STR was also modified to reduce the S+T% to 50%. Serines were changed by alanines trying to keep a maximum of 3 serines consecutives. Once designed, the new STRs were synthesized (GeneScript, plasmids pCM68-pCM70) and cloned in the pCM60 so the final sensors are Wsc1sp with modified STRs and fused to mNeonGreen (pCM71-73).

### Complementation assays

In *S. pombe*, deletion of both Mtl2 and Wsc1 mechanosensors is lethal (Cruz et al., 2013). For the functional complementation experiments, the strain sCM167 was transformed with the indicated plasmids with the Frozen-EZ transformation II kit (Zymo Research). These strains lack Wsc1sp (*wsc1Δ*) and Mtl2sp is under the control of the P81nmt1 repressive promoter and the sensor is expressed from the plasmid under the *wsc1sp* promoter. The strains were grown for 48h at 30°C in EMM-A agar plates supplemented with 5 µg/ml thiamine to repress the P81nmt1 promoter.

### Microscopy

Live-cell imaging experiments were performed on YE5S 2%-agarose pads or in micro-fabricated channels at room temperature (22-25°C). All images presented are single mid-sections obtained with an inverted spinning-disk confocal microscope equipped with a motorized stage and an automatic focus (Ti-Eclipse, Nikon, Japan), a CSUX1 spinning unit (Yokogawa) and a Prime BSI camera (Photometrics). Images were acquired with a 100X oil-immersion objective (CFI Plan Apo DM λ, Nikon), except for the cell-death quantification in channels, for which a 60X oil-immersion objective (CFI Apochromat λS, Nikon) was used. The microscope was operated with the Metamorph software (Molecular devices)

### Micro-fabricated channels

The design and fabrication of the flow channels was done as previously described (Haupt *et al*., 2018; Neeli-Venkata *et al*., 2021). The cells were loaded following an osmotic shock. The strains were grown on liquid YE5S at 25°C for 6 h, afterwards, 1 ml of the liquid culture was centrifugated at 3000 rpm 1 min, the supernatant discarded and the cells resuspended in 20µl of YE5S + 1M sorbitol. The cells were immediately loaded into the channels by applying pressure on the channel inlet with a syringe, usually yielding 1 cell per channel. The channels were washed two times and loaded with fresh YE5S, and incubated in a humid chamber overnight at 25°C. Next morning, 1 h before imaging at the microscope, the YE5S media was refreshed.

### Image analysis and quantifications

Local sensor enrichment was quantified using a semi-automated Matlab script (Mathworks) (Neeli-Venkata et al., 2021). The script removes the fluorescence background and allows the user to trace a line manually over the signal to be quantified, in this case the sensors-mNG signal between to cells that are compressed, based on their triangular shape (Contacts) and around cell tips with no contact (Free tip) (Figure S2). The line scan typically spans the site of contact and the cell sides with no contact along a distance of 10-14 µm. The fluorescence was averaged on a thickness of 12 px across the membrane, in order to average signals from both cells at sites of cell contacts, and fitted with a Gaussian, to extract basal fluorescent levels away from clusters, maximum signal in clusters, and width of the cluster. The ratio between the integral of the gaussian across its full width at mid height (FWMH), divided by the mean signal on cell sides multiplied by the FWMH, was used was used to compute the enrichment of the sensor. In order to compare the sensor enrichment at contacts and free tips, the script was modified in the latter cases to a thickness contour of 6 px, and the enrichment was multiplied by a factor 2 for normalization.

The signal intensity of the sensors at the cell periphery was measured in FIJI by drawing a segmented line of 5 px over the contour of the cell starting at the middle of one of the tips. The pixel intensity along the line was extracted from the profile and the background was removed, calculated from a 30×30 px square. The length was normalized by the total px length of the line and the intensity averaged on 50 bins of 0.02 length units (0-0.02 to 0.98-1). Tip was considered the bins 1-5, 21-30 and 46-50, and the sides the bins 6-20 and 31-45. The polarization ratio was calculated as the average signal intensity in the tips by the average intensity signal in the sides.

### Protein sequence similarity

We used the UniProt (The UniProt Consortium, 2025) align tool to analyze the similarity, in terms of identity, between the full protein sequences of the sensors and between the sensor’s domains as considered in table S3. The align tool uses Clustal Omega 1.2.4. For comparison between the full protein sequences of the sensors, the guide tree is shown, and for the domains the Percent Identity Matrix data was converted in a heat map.

### Statistical Analysis

All experiments in this study were repeated at least twice and quantifications performed in the number of cells or events specified in each figure. Graphs and statistical analyses were done in Prism 10 (GraphPad). Box plots show min to max values. To compare Contacts and Free tips for each strain, the non-parametric Mann-Whitney test was used. To compare the Contacts among different strains, the non-parametric Kruskal-Wallis test and Dunn’s test for multiple comparisons were used, in the compact letter display, strains under the same letter have similar medians and are significantly different to strains with other letters.

## ACKNOWLEDGMENTS

We thank Silke Hauf (Virginia tech) and Yolanda Sanchez (U. Salamanca) for sharing the material, as well as A. Boudaoud and S. Leon for discussion and careful reading of the manuscript. This work was supported by the Centre National de la Recherche Scientifique (CNRS), the Université Paris Cité, and grants from La Ligue Contre le Cancer (EL2021.LNCC/ NiM), and the Agence Nationale pour la Recherche (ANR, “CellWallSense” no. ANR-20-CE13-0003-02).

## Notes

### Competing Interest Statement

The authors have declared no competing interest.

