## Supplementary Material for "A CROSS-SPECIES ANALYSIS OF CELL WALL MECHANOSENSORS"

This supplementary material contains:

3 supplementary figures and figure legends and 4 supplementary tables

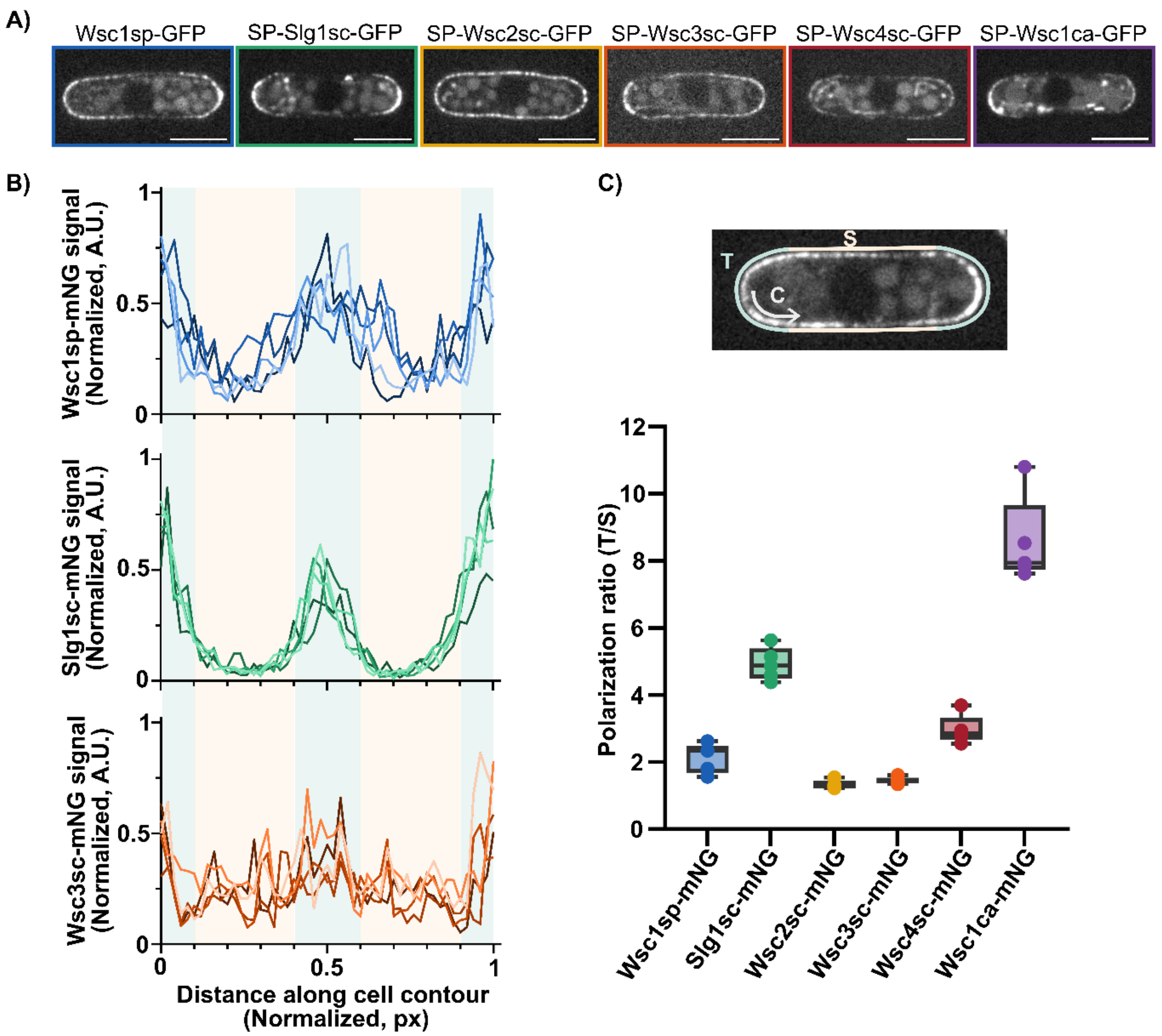

**Figure S1. Localization and polarization of foreign sensors**. **A)** Localization of the foreign sensors with their native signal peptide sequence. **B)** Signal intensity of the indicated sensors, with Wsc1sp signal peptide, along the cell contour. The green background corresponds to the tips and the yellow one to cell sides. Each line represents the contour of one cell (n=5 cells). **C)** The image of a representative cell shows where contour tracing starts, in green the region considered as the tips and in yellow the sides of the cell. The plot shows the polarization of the sensors tagged with mNeonGreen. This ratio is computed as the ratio of the signal intensity in the tip over the signal in the sides. Scale bars, 5 µm.

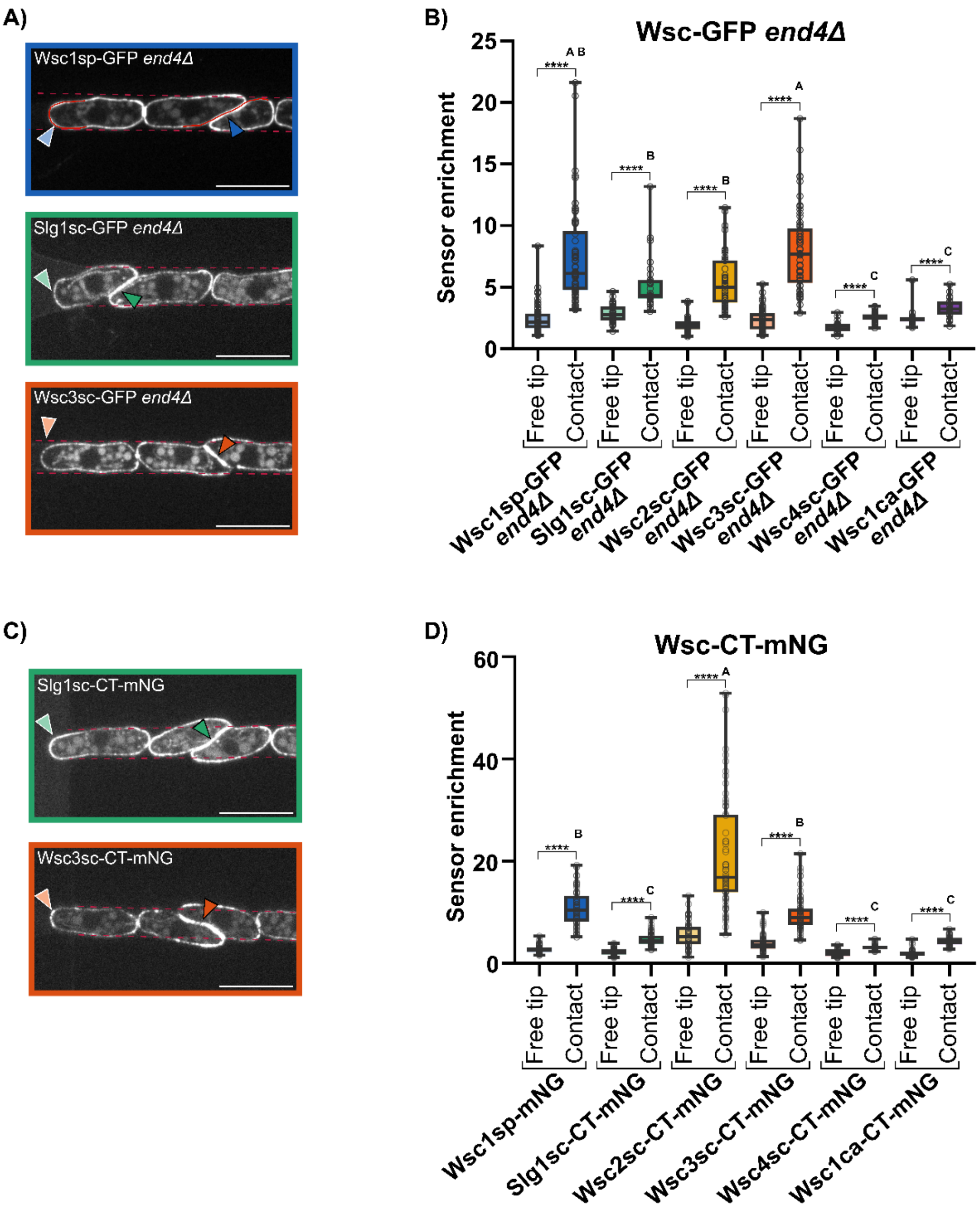
A)
Wsc1sp-GFP end4Δ
Slg1sc-GFP end4Δ
Wsc3sc-GFP end4Δ
B)
Wsc-GFP end4Δ
Sensor enrichment
25
20
15
10
5
0
****
A B
A
B
C
Free tip
Contact
Wsc1sp-GFP end4Δ
Slg1sc-GFP end4Δ
Wsc2sc-GFP end4Δ
Wsc3sc-GFP end4Δ
Wsc4sc-GFP end4Δ
Wsc1ca-GFP end4Δ
C)
Slg1sc-CT-mNG
Wsc3sc-CT-mNG
D)
Wsc-CT-mNG
Sensor enrichment
60
40
20
0
Wsc1sp-mNG
Slg1sc-CT-mNG
Wsc2sc-CT-mNG
Wsc3sc-CT-mNG
Wsc4sc-CT-mNG
Wsc1ca-CT-mNG

**Figure S2. Sensor localization and enrichment under mechanical stress in conditions when sensor endocytosis is perturbed**. **A) and C)** localization of the indicated sensors in microchannels. Light and dark-colored arrows point to examples of the quantified free tips and pressurized cell-cell contacts, respectively. The dashed red lines show the channels walls. The continuous red line shows examples of the line traced to compute the enrichment at cell tips and at cell-cell contacts. Scale bars, 10 µm. **B) and D)** Quantification of sensors enrichment at free tips and at cell contacts in an *end4Δ* mutant background and in sensors where the C-terminal tail has been swapped with that of Wsc1sp (Wsc-CT-mNG). In B) n= 66, 70, 39, 31, 45, 47, 57, 61, 29, 43, 25 and 27 cells in D) n = 54, 61, 59, 58, 61, 60, 58, 69, 29, 32, 48 and 53 cells. Comparison between free tips and contacts was done using the Mann-Whitney test, T-test P-values, ****, P < 0.0001. Scale bars, 10 µm.

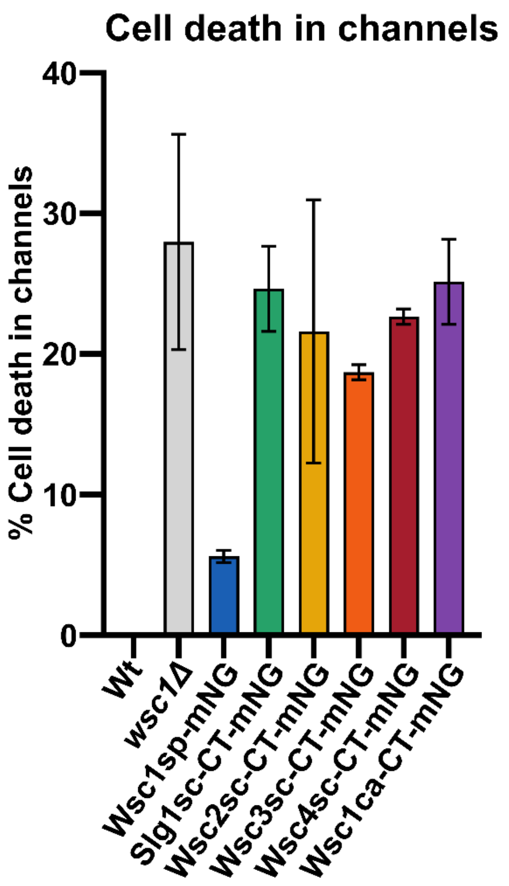

Cell death in channels
% Cell death in channels
40
30
20
10
0
Wt
wsc1Δ
Wsc1sp-mNG
Slg1sc-CT-mNG
Wsc2sc-CT-mNG
Wsc3sc-CT-mNG
Wsc4sc-CT-mNG
Wsc1ca-CT-mNG

**Figure S3. Cell death of cells expressing foreign sensors in microchannels.** Quantification of the percentage of cell death in microchannels for the Wsc-CT-mNG sensors. n = 170, 1216, 2227, 737, 1152, 1309, 955 and 977 cells.

**Table S1: Strains used in this study.** ALUH stand for the auxotrophies ade6M210, leu1-32, ura4D18 and his3D.

| Name | Genotype | Identifier | Source |
|---|---|---|---|
| Wild-type | *h+ ALU-* | AH146 | Minc lab stocks |
| Wild-type | *h- ALU-* | NM291 | Minc lab stocks |
| SP-SLG1sc-GFP | *h+ wsc1Δ::slg1sc-GFP:ura4+ ALU-* | CM7 | Integration of linear fragment pCM11 (oCM9+oCM10) in AH146 |
| SP-WSC2sc-GFP | *h+ wsc1Δ::wsc2sc-GFP:ura4+ ALU-* | CM8 | Integration of linear fragment pCM12 (oCM9+oCM10) in AH146 |
| SP-Wsc3sc-GFP | *h+ wsc1Δ::wsc3sc-GFP:ura4+ ALU-* | CM194 | Integration of linear fragment pCM13 (oCM9+oCM10) in AH146 |
| SP-WSC4sc-GFP | *h+ wsc1Δ::wsc4sc-GFP:ura4+ ALU-* | CM9 | Integration of linear fragment pCM14 (oCM9+oCM10) in AH146 |
| SP-WSC1ca-GFP | *h+ wsc1Δ::wsc1ca-GFP:ura4+ ALU-* | CM10 | Integration of linear fragment pCM15 (oCM9+oCM10) in AH146 |
| Slg1sc-mNG | *h+ wsc1Δ::slg1sc*$^{\Delta SP\text{-}wsc1spSP}$*-mNG:ura4+ ALU-* | CM101 | Integration of linear fragment from pCM52 in AH146. |
| Wsc2sc-mNG | *h+ wsc1Δ::wsc2sc*$^{\Delta SP\text{-}wsc1spSP}$*-mNG:ura4+ ALU-* | CM103 | Integration of linear fragment from pCM53 in AH146. |
| Wsc3sc-mNG | *h+ wsc1Δ::wsc3sc*$^{\Delta SP\text{-}wsc1spSP}$*-mNG:ura4+ ALU-* | CM105 | Integration of linear fragment from pCM54 in AH146. |
| Wsc4sc-mNG | *h+ wsc1Δ::wsc4sc*$^{\Delta SP\text{-}wsc1spSP}$*-mNG:ura4+ ALU-* | CM107 | Integration of linear fragment from pCM55 in AH146. |
| Wsc1ca-mNG | *h+ wsc1Δ::wsc1ca*$^{\Delta SP\text{-}wsc1spSP}$*-mNG:ura4+ ALU-* | CM109 | Integration of linear fragment from pCM56 in AH146. |
| *end4Δ* | *h- end4Δ::kanMX* | CM47 | Minc lab stocks |
| Wsc1sp-GFP_*end4Δ* | *h90 wsc1sp-GFP:ura4+ end4Δ::kanMX* | RN82 | Minc lab stocks (Neeli-Venkata et al., 2021) |
| SLG1sc-GFP_*end4Δ* | *wsc1Δ::slg1sc*$^{\Delta SP\text{-}wsc1spSP}$*-GFP:ura4+ end4::kanMX ALU-* | CM49 | Cross CM11 (*wsc1Δ::slg1sc*$^{\Delta SP\text{-}wsc1spSP}$*-GFP:ura4*) x CM47 |
| Wsc2sc-GFP_*end4Δ* | *wsc1Δ::wsc2sc*$^{\Delta SP\text{-}wsc1spSP}$*-GFP:ura4+ end4::kanMX ALU-* | CM51 | Cross CM13 (*wsc1Δ::wsc2sc*$^{\Delta SP\text{-}wsc1spSP}$*-GFP:ura4*) x CM47 |
| Wsc3sc-GFP_*end4Δ* | *wsc1Δ::wsc3sc*$^{\Delta SP\text{-}wsc1spSP}$*-GFP:ura4+ end4::kanMX ALU-* | CM53 | Cross CM15 (*wsc1Δ::wsc3sc*$^{\Delta SP\text{-}wsc1spSP}$*-GFP:ura4*) x CM47 |
| Wsc4sc-GFP_*end4Δ* | *wsc1Δ::wsc4sc*$^{\Delta SP\text{-}wsc1spSP}$*-GFP:ura4+ end4::kanMX ALU-* | CM55 | Cross CM17 (*wsc1Δ::wsc4sc*$^{\Delta SP\text{-}wsc1spSP}$*-GFP:ura4*) x CM47 |
| Wsc1ca-GFP_*end4Δ* | *wsc1Δ::wsc1ca*$^{\Delta SP\text{-}wsc1spSP}$*-GFP:ura4+ end4::kanMX ALU-* | CM57 | Cross CM19 (*wsc1Δ::wsc1ca*$^{\Delta SP\text{-}wsc1spSP}$*-GFP:ura4*) x CM47 |
| Slg1-CT-mNG | *h+ wsc1Δ::slg1sc*$^{\Delta SP\text{-}wsc1spSP\Delta CT\text{-}wsc1spCT}$*-GFP:ura4+ ALU-* | CM61 | Integration of linear fragment from pCM35 in AH146 |
| Wsc2-CT-mNG | *h+ wsc1Δ::wsc2sc*$^{\Delta SP\text{-}wsc1spSP\Delta CT\text{-}wsc1spCT}$*-GFP:ura4+ ALU-* | CM63 | Integration of linear fragment from pCM36 in AH146. |
| Wsc3-CT-mNG | *h+ wsc1Δ::wsc3sc*$^{\Delta SP\text{-}wsc1spSP\Delta CT\text{-}wsc1spCT}$*-GFP:ura4+ ALU-* | CM66 | Integration of linear fragment from pCM39 in AH146. |
| SPsp-Wsc4-CT-mNG | *h+ wsc1Δ::wsc4sc*$^{\Delta SP\text{-}wsc1spSP\Delta CT\text{-}wsc1spCT}$*-GFP:ura4+ ALU-* | CM68 | Integration of linear fragment from pCM38 in AH146. |
| Wsc1ca-CT-mNG | *h+ wsc1Δ::wsc4sc*$^{\Delta SP\text{-}wsc1spSP\Delta CT\text{-}wsc1spCT}$*-GFP:ura4+ ALU-* | CM70 | Integration of linear fragment from pCM37 in AH146. |
| Slg1scWSC-mNG | *h+ wsc1*$^{\Delta WSC\text{-}slg1scWSC}$*-mNG:ura4+ ALU-* | CM85 | Integration of linear fragment from pCM44 (Ppu21) in AH146 |
| Wsc3scWSC-mNG | *h+ wsc1*$^{\Delta WSC\text{-}wsc3scWSC}$*-mNG:ura4+ ALU-* | CM92 | Integration of linear fragment from pCM46 (Ppu21) in AH146 |
| Wsc2scWSC-mNG | *h+ wsc1*$^{\Delta WSC\text{-}wsc2scWSC}$*-mNG:ura4+ ALU-* | CM111 | Integration of linear fragment from pCM57 (Ppu21) in AH146 |
| Wsc4scWSC-mNG | *h+ wsc1*$^{\Delta WSC\text{-}wsc4scWSC}$*-mNG:ura4+ ALU-* | CM113 | Integration of linear fragment from pCM58 (Ppu21) in AH146 |
| Wsc1caWSC-mNG | *h+ wsc1*$^{\Delta WSC\text{-}wsc1caWSC}$*-mNG:ura4+ ALU-* | CM115 | Integration of pCM59 linear fragment (Eco721+KpnI) in AH146 |

| | | | |
|---|---|---|---|
| Slg1scSTR-mNG | *h- wsc1*$^{\Delta STR\text{-}slg1scSTR}$*-mNG:ura4+ ALU-* | CM123 | Integration of linear fragment from pCM63 in NM291 |
| Wsc2scSTR-mNG | *h- wsc1*$^{\Delta STR\text{-}wsc2scSTR}$*-mNG:ura4+ ALU-* | CM125 | Integration of linear fragment from pCM64 in NM291 |
| Wsc3scSTR-mNG | *h- wsc1*$^{\Delta STR\text{-}wsc3scSTR}$*-mNG:ura4+ ALU-* | CM127 | Integration of linear fragment from pCM65 in NM291 |
| Wsc4scSTR-mNG | *h- wsc1*$^{\Delta STR\text{-}wsc4scSTR}$*-mNG:ura4+ ALU-* | CM129 | Integration of linear fragment from pCM66 in NM291 |
| Wsc1caSTR-mNG | *h- wsc1*$^{\Delta STR\text{-}wsc1caSTR}$*-mNG:ura4+ ALU-* | CM131 | Integration of linear fragment from pCM67 in NM291 |
| Wsc1-mNeonGreen | *h+ wsc1-mNG:ura4+ ALU-* | CM151 | Integration of linear fragment from pCM60 (ppU21I) in AH146 |
| Wsc4scSTRshort-mNG | *h+ wsc1*$^{\Delta STR\text{-}wsc4scSTRshort}$*-mNG:ura4+ ALU-* | CM153 | Integration of linear fragment from pCM67 (ppU21I) in AH146 |
| Wsc1caSTRlong-mNG | *h+ wsc1*$^{\Delta STR\text{-}wsc1caSTRlong}$*-mNG:ura4+ ALU-* | CM155 | Integration of linear fragment from pCM66 (ppU21I) in AH146 |
| *kin1Δ* | *h+ kin1Δ::kanMX ALU-* | AH139 | Minc lab stocks |
| Wsc1sp-mNG_*kin1Δ* | *h- wsc1-mNG:ura4+ kin1Δ::kanMX ALU-* | CM157 | Cross CM149 (*h- kin1Δ::kanMX*) x CM151 |
| Wsc4scSTR-mNG_*kin1Δ* | *h+ wsc1*$^{\Delta STR\text{-}wsc4scSTR}$*-mNG:ura4+ kin1Δ::kanMX ALU-* | CM159 | Cross AH139 x CM129 |
| Wsc1caSTR-mNG_*kin1Δ* | *h+ wsc1sp*$^{\Delta STR\text{-}wsc1caSTR}$*-mNG:ura4+ kin1Δ::kanMX ALU-* | CM161 | Cross AH139 x CM131 |
| Wsc1spSTR50-mNG | *h+ wsc1*$^{\Delta STR\text{-}wsc1spSTR50}$*-mNG:ura4+ ALU-* | CM197 | Integration of linear fragment from pCM71 (ppU21I) in AH146 |
| Wsc4scSTR76-mNG | *h+ wsc1*$^{\Delta STR\text{-}wsc4scSTR76}$*-mNG:ura4+ ALU-* | CM189 | Integration of linear fragment from pCM72 (ppU21I) in AH146 |
| Wsc1caSTR76-mNG | *h+ wsc1*$^{\Delta STR\text{-}wsc1scSTR76}$*-mNG:ura4+ ALU-* | CM191 | Integration of linear fragment from pCM73 (ppU21I) in AH146 |
| *wsc1Δ* | *h- wsc1Δ::kanMX ALUH-* | GRG15 | Cruz et al., 2013 |
| *wsc1Δ* | *h- wsc1Δ::natMX ALUH-* | CM163 | Marker switch, kanMX by natMX in strain GRG15 |
| *wsc1Δ*_mtl2 switch-off | *h+ wsc1Δ::natMX kan-81xnmt1-mtl2*$^{+}$ *ALUH-* | CM167 | Cross CM163 x SC80 (Cruz et al., 2013) |
| *wsc1Δ*_mtl2 switch-off_Slg1sc-mNG | *h+ wsc1Δ::natMX kan-81xnmt1-mtl2*$^{+}$ *+ plasmid pCM52 ALUH-* | CM168 | Plasmid pCM52 transformation in CM167 strain. |
| *wsc1Δ*_mtl2 switch-off_Wsc2sc-mNG | *h+ wsc1Δ::natMX kan-81xnmt1-mtl2*$^{+}$ *+ plasmid pCM53 ALUH-* | CM169 | Plasmid pCM53 transformation in CM167 strain. |
| *wsc1Δ*_mtl2 switch-off_Wsc3sc-mNG | *h+ wsc1Δ::natMX kan-81xnmt1-mtl2*$^{+}$ *+ plasmid pCM54 ALUH-* | CM170 | Plasmid pCM54 transformation in CM167 strain. |
| *wsc1Δ*_mtl2 switch-off_Wsc4sc-mNG | *h+ wsc1Δ::natMX kan-81xnmt1-mtl2*$^{+}$ *+ plasmid pCM55 ALUH-* | CM171 | Plasmid pCM55 transformation in CM167 strain. |
| *wsc1Δ*_mtl2 switch-off_Wsc1ca-mNG | *h+ wsc1Δ::natMX kan-81xnmt1-mtl2*$^{+}$ *+ plasmid pCM56 ALUH-* | CM172 | Plasmid pCM56 transformation in CM167 strain. |
| *wsc1Δ*_mtl2 switch-off_Slg1sc-CT-mNG | *h+ wsc1Δ::natMX kan-81xnmt1-mtl2*$^{+}$ *+ plasmid pCM35 ALUH-* | CM173 | Plasmid pCM35 transformation in CM167 strain. |
| *wsc1Δ*_mtl2 switch-off_Wsc2sc-CT-mNG | *h+ wsc1Δ::natMX kan-81xnmt1-mtl2*$^{+}$ *+ plasmid pCM36 ALUH-* | CM174 | Plasmid pCM36 transformation in CM167 strain. |
| *wsc1Δ*_mtl2 switch-off_Wsc3sc-CT-mNG | *h+ wsc1Δ::natMX kan-81xnmt1-mtl2*$^{+}$ *+ plasmid pCM37 ALUH-* | CM175 | Plasmid pCM37 transformation in CM167 strain. |

| | | | |
|---|---|---|---|
| *wsc1Δ*_mtl2 switch-off_Wsc4sc-CT-mNG | *h+ wsc1Δ::natMX kan-81xnmt1-mtl2*$^{+}$ + *plasmid pCM38 ALUH-* | CM176 | Plasmid pCM38 transformation in CM167 strain. |
| *wsc1Δ*_mtl2 switch-off_Wsc1ca-CT-mNG | *h+ wsc1Δ::natMX kan-81xnmt1-mtl2*$^{+}$ + *plasmid pCM39 ALUH-* | CM177 | Plasmid pCM39 transformation in CM167 strain. |
| *wsc1Δ*_mtl2 switch-off_Slg1sc-WSC-mNG | *h+ wsc1Δ::natMX kan-81xnmt1-mtl2*$^{+}$ + *plasmid pCM44 ALUH-* | CM178 | Plasmid pCM44 transformation in CM167 strain. |
| *wsc1Δ*_mtl2 switch-off_Wsc2sc-WSC-mNG | *h+ wsc1Δ::natMX kan-81xnmt1-mtl2*$^{+}$ + *plasmid pCM57 ALUH-* | CM179 | Plasmid pCM57 transformation in CM167 strain. |
| *wsc1Δ*_mtl2 switch-off_Wsc3sc-WSC-mNG | *h+ wsc1Δ::natMX kan-81xnmt1-mtl2*$^{+}$ + *plasmid pCM46 ALUH-* | CM180 | Plasmid pCM46 transformation in CM167 strain. |
| *wsc1Δ*_mtl2 switch-off_Wsc4sc-WSC-mNG | *h+ wsc1Δ::natMX kan-81xnmt1-mtl2*$^{+}$ + *plasmid pCM58 ALUH-* | CM181 | Plasmid pCM58 transformation in CM167 strain. |
| *wsc1Δ*_mtl2 switch-off_Wsc1ca-WSC-mNG | *h+ wsc1Δ::natMX kan-81xnmt1-mtl2*$^{+}$ + *plasmid pCM59 ALUH-* | CM182 | Plasmid pCM59 transformation in CM167 strain. |
| *wsc1Δ*_mtl2 switch-off_Slg1sc-STR-mNG | *h+ wsc1Δ::natMX kan-81xnmt1-mtl2*$^{+}$ + *plasmid pCM61 ALUH-* | CM183 | Plasmid pCM61 transformation in CM167 strain. |
| *wsc1Δ*_mtl2 switch-off_Wsc2sc-STR-mNG | *h+ wsc1Δ::natMX kan-81xnmt1-mtl2*$^{+}$ + *plasmid pCM62 ALUH-* | CM184 | Plasmid pCM62 transformation in CM167 strain. |
| *wsc1Δ*_mtl2 switch-off_Wsc3sc-STR-mNG | *h+ wsc1Δ::natMX kan-81xnmt1-mtl2*$^{+}$ + *plasmid pCM63 ALUH-* | CM185 | Plasmid pCM63 transformation in CM167 strain. |
| *wsc1Δ*_mtl2 switch-off_Wsc4sc-STR-mNG | *h+ wsc1Δ::natMX kan-81xnmt1-mtl2*$^{+}$ + *plasmid pCM64 ALUH-* | CM186 | Plasmid pCM64 transformation in CM167 strain. |
| *wsc1Δ*_mtl2 switch-off_Wsc1ca-STR-mNG | *h+ wsc1Δ::natMX kan-81xnmt1-mtl2*$^{+}$ + *plasmid pCM65 ALUH-* | CM187 | Plasmid pCM65 transformation in CM167 strain. |
| *wsc1Δ*_mtl2 switch-off_Wsc1sp-mNG | *h+ wsc1Δ::natMX kan-81xnmt1-mtl2*$^{+}$ + *plasmid pCM60 ALUH-* | CM188 | Plasmid pCM60 transformation in CM167 strain, |

### Table S2: Oligonucleotides used in this study

| Identifier | Name | Sequence |
|---|---|---|
| oCM9 | wsc_trans_F | TGTTCGCCTAGGTTCGACTG |
| oCM10 | wsc_trans_R | CAAGCACCAATCGACCAGGA |
| oCM11 | SP_SLG1sc_V_R | GAAAAGCAGTTGACGTACTCGTCAGCCGCCACCAACC |
| oCM12 | SP_SLG1sc_F_F | CTCGGTTGGTGGCGGCTGACGAGTACGTCAACTGCTTTTCTTCATTACC |
| oCM13 | SP_WSC2sc_V_R | CAAGCCTTATAGGTAAATTGGTCAGCCGCCACCAACC |
| oCM14 | SP_WSC2sc_F_F | GTTGGTGGCGGCTGACCAATTTACCTATAAGGCTTGCTACTCTGC |
| oCM15 | SP_WSC3sc_V_R | AGCAACCCTCATAGTTAAAGTCAGCCGCCACCAACC |
| oCM16 | SP_WSC3sc_F_F | GGTTGGTGGCGGCTGACTTTAACTATGAGGGTTGCTATTCAGCTG |
| oCM17 | SP_WSC4sc_V_R | TGAGAAGAACAGACAGATTGGTCAGCCGCCACCAACC |
| oCM18 | SP_WSC4sc_F_F | CTCGGTTGGTGGCGGCTGACCAATCTGTCTGTTCTTCTCAAAACACTGC |
| oCM19 | SP_WSC1ca_V_R | GAGGCTGGTGCAGTATAATCGTCAGCCGCCACCAACC |
| oCM20 | SP_WSC1ca_F_F | CTCGGTTGGTGGCGGCTGACGATTATACTGCACCAGCCTCAAATCTTGG |
| oCM21 | SP_Vector_F | GCATGGATGAACTATACAAAGGCCGCAACTAAAACAACATTTTTCTTT |
| oCM22 | SP_Fragment_R | ATGTTGTTTTAGTTGCGGCCTTTGTATAGTTCATCCATGCCATGTGT |
| oCM39 | CTailswap_Slg1_CT_F | ATTTTGTTGATTGTATATTTCAGGAGATTCAAAATTAGAATGAGCGACTC |
| oCM40 | CTailswap_Slg1_vector_R | GAATCTCCTGAAATATACAATCAACAAAATGCAAAGAGCAATAGC |
| oCM41 | CTailswap_Wsc2_CT_F | TTGTTCTTCTTTGTGTATTTCAGGAGATTCAAAATTAGAATGAGCGACTC |
| oCM42 | CTailswap_Wsc2_vector_R | GAATCTCCTGAAATACACAAAGAAGAACAAGGCAAGAGC |
| oCM43 | CTailswap_Wsc3_CT_F | TTATTCTTGATCTGGTATTTCAGGAGATTCAAAATTAGAATGAGCGACTC |
| oCM44 | CTailswap_Wsc3_vector_R | GAATCTCCTGAAATACCAGATCAAGAATAATAGAATAAGTATTATAAAGATTACGCCAAAG |
| oCM45 | CTailswap_Wsc4_CT_F | CTTATCTACTTAATTTATTTCAGGAGATTCAAAATTAGAATGAGCGACTC |
| oCM46 | CTailswap_Wsc4_vector_R | GAATCTCCTGAAATAAATTAAGTAGATAAGAATGCATATAATCACTAAGCAGACAAC |
| oCM47 | CTailswap_Wsc1ca_CT_F | GGTTTCTTTTATATGTATTTCAGGAGATTCAAAATTAGAATGAGCGACTC |
| oCM48 | CTailswap_Wsc1ca_vector_R | GAATCTCCTGAAATACATATAAAAGAAACCTCCAGCCACTAATAGGA |
| oCM49 | CTailswap_NG_F | CGTGTCACAAATTTGCGGATCCCCGGGTTAATTAACG |
| oCM50 | CTailswap_CT_R | TAACCCGGGGATCCGCAAATTTGTGACACGCAATATCTTTCTCG |
| oCM51 | CTailswap_vector_F | GAACTTTACAAATAGAACAACATTTTTCTTTTTTACTTTATTTAAATTCAACCAAAAAAT |
| oCM52 | CTailswap_NG_R | AAGAAAAATGTTGTTCTATTTGTAAAGTTCATCCATACCCATAACATCAGT |
| oCM68 | Wsc1sp_Slg1scWSC_F_F | TATCAGTTAGATAATGGGGTTTcGCAAACCACG |
| oCM69 | Wsc1sp_Slg1scWSC_F_R | ATCTCCTGAAATAGAAGTATAGGAAAATGCACAATG |
| oCM70 | Wsc1sp_Slg1scWSC_V_F | TTCCTATACTTCTATTTCAGGAGATTCAAAATTAGAATGAGCGACT |
| oCM71 | Wsc1sp_Slg1scWSC_V_R | GCgAAACCCCATTATCTAACTGATAAACGCTATAGGCATCTTCG |
| oCM72 | Wsc1sp_Wsc3scWSC_F_F | CGTTAACGCCAATGGGGTTTcGCAAACCAC |
| oCM73 | Wsc1sp_Wsc3scWSC_F_R | TCTCCTGAAATAGAAGTATAGGAAAATGCACAATG |
| oCM74 | Wsc1sp_Wsc3scWSC_V_F | TCCTATACTTCTATTTCAGGAGATTCAAAATTAGAATGAGCGACT |

| | | |
|---|---|---|
| oCM75 | Wsc1sp_Wsc3scWSC_V_R | AAACCCCATTGGCGTTAACGTATACGTTCATATATGAAGAAC |
| oCM83 | 5UTRg2_Wsc1_F | CGTGGGTACTTCGACATGGT |
| oCM92 | mNG_V_F | CGGATCCCCGGGTTAATTAACG |
| oCM93 | mNG_V_R | GTCAGCCGCCACCAACC |
| oCM94 | Slg1_mNG_F_F | TTGGTGGCGGCTGACGAGTACGTCAAC |
| oCM95 | Slg1_mNG_F_R | TAACCCGGGGATCCGATCAGCTTCATCAGGATTTACCACAG |
| oCM96 | Wsc2_mNG_F_F | TTGGTGGCGGCTGACCAATTTACCTAT |
| oCM97 | Wsc2_mNG_F_R | TAACCCGGGGATCCGACGAAGTGAAGAGTTGTTTGAATCTTTGG |
| oCM98 | Wsc3-mNG_F_F | TTGGTGGCGGCTGACTTTAACTATGAGG |
| oCM99 | Wsc3-mNG_F_R | TAACCCGGGGATCCGAGCACGATTGTGTGAAACAGTTGA |
| oCM100 | Wsc4-mNG_F_F | TTGGTGGCGGCTGACCAATCTGTCTGT |
| oCM101 | Wsc4-mNG_F_R | TAACCCGGGGATCCGTTCATTCATAAGGCGCAAAACGCGT |
| oCM102 | Wsc1ca-mNG_F_F | TTGGTGGCGGCTGACGATTATACTGCA |
| oCM103 | Wsc1ca-mNG_F_R | TAACCCGGGGATCCGTTTTGTGTCGTCTGGATTGGCAACT |
| oCM104 | WSC_V_F | AATGGGGTTTcGCAAACCACG |
| oCM105 | WSC_V_R | GTCAGCCGCCACCAACC |
| oCM106 | WSC_wsc2_F_F | TTGGTGGCGGCTGACCAATTTACCTAT |
| oCM107 | WSC_wsc2_F_R | TTTGCgAAACCCCATTATTGTTGATATAGACATTCATAGCGGATGATCC |
| oCM108 | WSC_wsc4_F_F | TTGGTGGCGGCTGACCAATCTGTCTGT |
| oCM109 | WSC_wsc4_F_R | TTTGCgAAACCCCATTTAAGTAAATGTAACCAAACAAATCTTTGTCCGCA |
| oCM110 | WSC_wsc1ca_F_F | TTGGTGGCGGCTGACGATTATACTGCA |
| oCM111 | WSC_wsc1ca_F_R | TTTGCgAAACCCCATTTAAGCCTTTGTAAATGTTATAAGCATTAGATCCACC |
| oCM112 | Wsc1sp_mNG_V_F | CAAATTTGCGGATCCCCGGG |
| oCM113 | Wsc1sp_mNG_V_R | TTAAAAAGACCATGGAAATTAAATCGTGCGT |
| oCM114 | Wsc1sp_mNG_F_F | GATTTAATTTCCATGGTCTTTTTAAATTCCTCTCCC |
| oCM115 | Wsc1sp_mNG_F_R | GGGGATCCGCAAATTTGTGACACGCAATATCTT |
| oCM116 | STR_V_F | TCAAACCATACGTCCTTGAATGCAG |
| oCM117 | STR_V_R | AACCGTGGTTTGCgAAACC |
| oCM118 | Slg1_STR_F_F | TcGCAAACCACGGTTTCAGATACCAATTCGAATAGCATTTCTTCTTCAG |
| oCM119 | Slg1_STR_F_R | CAAGGACGTATGGTTTGATGCTCCAACATTGGCTTTCTTCT |
| oCM120 | Wsc2_STR_F_F | TcGCAAACCACGGTTGCTGCATCTACAGCGGATTCA |
| oCM121 | Wsc2_STR_F_R | CAAGGACGTATGGTTTGAACCAGCTATCGCGCCAC |
| oCM122 | Wsc3_STR_F_F | TcGCAAACCACGGTTGAAACATTTGTCTCCAGTGTAGAATCGTCA |
| oCM123 | Wsc3_STR_F_R | CAAGGACGTATGGTTTGAACCAGCAATTGCTCCTCCAG |
| oCM124 | Wsc4_STR_F_F | TcGCAAACCACGGTTGGGCAAACTCCCTTATCGTCAGT |
| oCM125 | Wsc4_STR_F_R | CAAGGACGTATGGTTTGAAGCTATTTTGCCAGGACTATCCC |
| oCM126 | Wsc1ca_STR_F_F | TcGCAAACCACGGTTGCTTCTGCTAGTTCCAGTTCATCAT |
| oCM127 | Wsc1ca_STR_F_R | CAAGGACGTATGGTTTGATGCTCCAACTGAAGATGACTTCTTCT |
| oCM134 | oCM134_Ws1caSTRlong_F | GCCAGTGCAACAACTTCTAGTGGAAATAATTCCGACGACAGTGATCAGTCTAC |
| oCM135 | oCM135_Wsc1caSTRlong_R | TCCACTAGAAGTTGTTGCACTGGC |
| oCM136 | oCM136_Wsc4STRShort-mNG_F | ACAATGGACACTAATATATCAGAGATTACTTCACG |
| oCM137 | oCM137_Wsc4STRShort-mNG_R | TTTGGTGGAGGTAGTAGTCGGTAATG |
| oCM143 | CM143_Wsc1spSTR50co_F_F | TcGCAAACCACGGTTTCATCGAGCTCCGTTAGTGCTAC |
| oCM144 | CM144_Wsc1spSTR50co_F_R | CAAGGACGTATGGTTTGAAGCTTTAGACGCGTTTAAGGCT |

#### Table S3: Sensor domain lengths

| Sensor | Signal peptide | WSC domain | STR domain | Transmembrane domain | Cytoplasmic tail | Total length (aa) |
|---|---|---|---|---|---|---|
| Slg1sc | 1-21<br>(21 aa) | 22-110<br>(89 aa) | 111-264<br>(154 aa) | 265-285<br>(21 aa) | 286-378<br>(93 aa) | 378 |
| Wsc2sc | 1-23<br>(23 aa) | 24-118<br>(95 aa) | 119-325<br>(207 aa) | 326-346<br>(21 aa) | 347-503<br>(157 aa) | 503 |
| Wsc3sc | 1-38<br>(38 aa) | 39-132<br>(94 aa) | 133-384<br>(252 aa) | 385-405<br>(21 aa) | 406-556<br>(151 aa) | 556 |
| Wsc4sc | 1-26<br>(26 aa) | 27-110<br>(84 aa) | 111-414<br>(304 aa) | 415-435<br>(21 aa) | 436-605<br>(170 aa) | 605 |
| Wsc1ca | 1-23<br>(23 aa) | 24-116<br>(93 aa) | 117-231<br>(115 aa) | 232-253<br>(22 aa) | 254-358<br>(105 aa) | 358 |
| Wsc1sp | 1-29<br>(29 aa) | 30-119<br>(90 aa) | 120-292<br>(173 aa) | 293-315<br>(23 aa) | 316-374<br>(59 aa) | 374 |

### Table S4: Plasmids used in this study

| Identifier | Name | Source |
|---|---|---|
| pRZ21 | pAL-wsc1-GFP:ura4+ | Cruz et al., 2013 |
| pCM29 | pFA6a_NeonGreen_kan | Silke Hauf |
| pCM1 | Slg1sc | This study. Eurofins Genomics gene synthesis |
| pCM2 | Wsc2sc | This study. Eurofins Genomics gene synthesis |
| pCM3 | Wsc3sc | This study. Eurofins Genomics gene synthesis |
| pCM4 | Wsc4sc | This study. Eurofins Genomics gene synthesis |
| pCM5 | Wsc1ca | This study. Eurofins Genomics gene synthesis |
| pCM10 | pRZ21_modified | This study. Site directed mutagenesis on pRZ21 |
| pCM11 | Slg1sc-GFP | This study. Insertion of XbaI-Notl fragment from pCM1 into NheI-Notl sites of pCM10 |
| pCM12 | Wsc2sc-GFP | This study. Insertion of XbaI-Notl fragment from pCM2 into NheI-Notl sites of pCM10 |
| pCM13 | Wsc3sc-GFP | This study. Insertion of XbaI-Notl fragment from pCM3 into NheI-Notl sites of pCM10 |
| pCM14 | Wsc4sc-GFP | This study. Insertion of XbaI-Notl fragment from pCM4 into NheI-Notl sites of pCM10 |
| pCM15 | Wcs1ca-GFP | This study. Insertion of XbaI-Notl fragment from pCM5 into NheI-Notl sites of pCM10 |
| pCM16 | SPsp-Slg1sc-GFP | This study. Gibson assembly fragments oCM11+oCM21 (pRZ21) and oCM12+oCM22 (pCM11) |
| pCM17 | SPsp-Wsc2sc-GFP | This study. Gibson assembly fragments oCM13+oCM21 (pRZ21) and oCM14+oCM22 (pCM12) |
| pCM18 | SPsp-Wsc3sc-GFP | This study. Gibson assembly fragments oCM15+oCM21 (pRZ21) and oCM16+oCM22 (pCM13) |
| pCM19 | SPsp-Wsc4sc-GFP | This study. Gibson assembly fragments oCM17+oCM21 (pRZ21) and oCM18+oCM22 (pCM14) |
| pCM20 | SPsp-WSsc1ca-GFP | This study. Gibson assembly fragments oCM19+oCM21 (pRZ21) and oCM20+oCM22 (pCM15) |
| pCM52 | SPsp-Slg1sc-mNG | This study. Gibson assembly fragments oCM94+oCM95 (pCM16) and oCM92+oCM93 (pCM35) |
| pCM53 | SPsp-Wsc2sc-mNG | This study. Gibson assembly fragments oCM96+oCM97 (pCM17) and oCM92+oCM93 (pCM35) |
| pCM54 | SPsp-Wsc3sc-mNG | This study. Gibson assembly fragments oCM98+oCM99 (pCM18) and oCM92+oCM93 (pCM35) |
| pCM55 | SPsp-Wsc4sc-mNG | This study. Gibson assembly fragments oCM100+oCM101 (pCM19) and oCM92+oCM93 (pCM35) |
| pCM56 | SPsp-Wsc1ca-mNG | This study. Gibson assembly fragments oCM102+oCM103 (pCM20) and oCM92+oCM93 (pCM35) |
| pCM60 | Wsc1sp-mNG | This study. Gibson assembly fragments oCM112+oCM113 (pCM29) and oCM114+oCM115 (PRZ21) |
| pCM35 | SPsp-Slg1sc-CTsp-mNG | This study. Gibson assembly fragments oCM40+oCM51 (pCM16),oCM39+oCM50 (pRZ21) and oCM49+oCM52 (pCM29) |
| pCM36 | SPsp-Wsc2sc-CTsp-mNG | This study. Gibson assembly fragments oCM42+oCM51 (pCM17),oCM41+oCM50 (pRZ21) and oCM49+oCM52 (pCM29) |
| pCM37 | SPsp-Wsc1ca-CTsp-mNG | This study. Gibson assembly fragments oCM48+oCM51 (pCM20),oCM47+oCM50 (pRZ21) and oCM49+oCM52 (pCM29) |
| pCM38 | SPsp-Wsc4sc-CTsp-mNG | This study. Gibson assembly fragments oCM46+oCM51 (pCM19),oCM45+oCM50 (pRZ21) and oCM49+oCM52 (pCM29) |
| pCM39 | SPsp-Wsc3sc-CTsp-mNG | This study. Gibson assembly fragments oCM44+oCM51 (pCM18),oCM43+oCM50 (pRZ21) and oCM49+oCM52 (pCM29) |
| pCM44 | Wsc1sp-Slg1scWSC-mNG | This study. Gibson assembly fragments oCM68+oCM69 (pRZ21) and oCM70+oCM71 (pCM35) |
| pCM46 | Wsc1sp-Wsc3scWSC-mNG | This study. Gibson assembly fragments oCM72+oCM73 (pRZ21) and oCM74+oCM75 (pCM39) |
| pCM57 | Wsc1sp-Wsc2scWSC-mNG | This study. Gibson assembly fragments oCM106+oCM107 (pCM17) and oCM104+oCM105 (pCM44) |
| pCM58 | Wsc1sp-Wsc4scWSC-mNG | This study. Gibson assembly fragments oCM108+oCM109 (pCM19) and oCM104+oCM105 (pCM44) |

| | | |
|---|---|---|
| pCM59 | Wsc1sp-Wsc1caWSC-mNG | This study. Gibson assembly fragments oCM110+oCM111 (pCM20) and oCM104+oCM105 (pCM44) |
| pCM61 | Wsc1sp-Slg1scSTR-mNG | This study. Gibson assembly fragments oCM116+oCM117 (pCM60) and oCM118+oCM119 (pCM52) |
| pCM62 | Wsc1sp-Wsc2scSTR-mNG | This study. Gibson assembly fragments oCM116+oCM117 (pCM60) and oCM120+oCM121 (pCM53) |
| pCM63 | Wsc1sp-Wsc3scSTR-mNG | This study. Gibson assembly fragments oCM116+oCM117 (pCM60) and oCM122+oCM123 (pCM54) |
| pCM64 | Wsc1sp-Wsc4scSTR-mNG | This study. Gibson assembly fragments oCM116+oCM117 (pCM60) and oCM124+oCM125 (pCM55) |
| pCM65 | Wsc1sp-Wsc1caSTR-mNG | This study. Gibson assembly fragments oCM116+oCM117 (pCM60) and oCM126+oCM127 (pCM56) |
| pCM67 | Wsc1sp-Wsc4-STRshort-mNG | This study. Circularization fragment with phosphorilated oligos oCM136+oCM137 (pCM64) |
| pCM66 | Wsc1sp-Wsc1caSTRlong-mNG | This study. Gibson assembly fragments oCM135+oCM51 (pCM65) and oCM134+oCM52 (pCM65) |
| pCM68 | Wsc1sp-STR50co | This study. Gene synthesis by GeneScript |
| pCM69 | Wsc4sc-STR76 | This study. Gene synthesis by GeneScript |
| pCM70 | Wsc1ca-STR76 | This study. Gene synthesis by GeneScript |
| pCM71 | Wsc1sp-STR50-mNG | This study. Gibson assembly fragments oCM116+oCM117 (pCM60) and oCM143+oCM144 (pCM68) |
| pCM72 | Wsc1sp-Wsc4scSTR76-mNG | This study. Gibson assembly fragments oCM116+oCM117 (pCM60) and oCM124+oCM125 (pCM69) |
| pCM73 | Wsc1sp-Wsc1caSTR76-mNG | This study. Gibson assembly fragments oCM116+oCM117 (pCM60) and oCM126+oCM127 (pCM70) |
